# Cytotoxic rhamnolipid micelles drive acute virulence in *Pseudomonas aeruginosa*

**DOI:** 10.1101/2023.10.13.562257

**Authors:** Qi Xu, Donghoon Kang, Matthew D. Meyer, Christopher L. Pennington, Citrupa Gopal, Jeffrey W. Schertzer, Natalia V. Kirienko

**Author notes:** Address correspondence to Natalia V. Kirienko.

## Abstract

*Pseudomonas aeruginosa* is an opportunistic human pathogen that has developed multi- or even pan-drug resistance towards most frontline and last resort antibiotics, leading to increasing infections and deaths among hospitalized patients, especially those with compromised immune systems. Further complicating treatment, *P. aeruginosa* produces numerous virulence factors that contribute to host tissue damage and immune evasion, promoting bacterial colonization and pathogenesis. In this study, we demonstrate the importance of rhamnolipid production in host-pathogen interactions. Secreted rhamnolipids form micelles that exhibited highly acute toxicity towards murine macrophages, rupturing the plasma membrane and causing organellar membrane damage within minutes of exposure. While rhamnolipid micelles (RMs) were particularly toxic to macrophages, they also caused membrane damage in human lung epithelial cells, red blood cells, Gram-positive bacteria, and even non-cellular models like giant plasma membrane vesicles. Most importantly, rhamnolipid production strongly correlated to *P. aeruginosa* virulence against murine macrophages in various panels of clinical isolates. Altogether, our findings suggest that rhamnolipid micelles are highly cytotoxic virulence factors that drive acute cellular damage and immune evasion during *P. aeruginosa* infections.

## Introduction

*Pseudomonas aeruginosa* is a Gram-negative opportunistic human pathogen that causes a variety of nosocomial infections, including ventilator-associated pneumonia, urinary tract infections, and soft tissue infections.(1, 2) It is also one of the most common causes of secondary bacterial infection in influenza and COVID-19 patients.(3) Unfortunately, *P. aeruginosa* infections are becoming increasingly difficult to treat due to the pathogen’s resistance to many front-line antibiotics and its ability to evade and subvert host immune responses.(4, 5) During acute infection, *P. aeruginosa* notoriously secretes a panoply of virulence factors like the elastases LasA and LasB and proteases AprA and PrpL that degrade host immune proteins such as immunoglobin G and complement factors.(5–9) *P. aeruginosa* also produces various toxins, such as those secreted by the type III secretion system, that damage and kill phagocytic immune cells.(10, 11) We recently demonstrated that production of the siderophore pyoverdine was important for *P. aeruginosa* virulence against murine macrophages.(12) During chronic infections, particularly in the lungs of cystic fibrosis patients*, P. aeruginosa* forms dense biofilms – bacterial communities reversibly attached to host tissue through the secretion of adhesion proteins, extracellular DNA, and exopolysaccharides – that are impervious to host immune cells and antimicrobial therapy.(13–15)

While most pathogen immune evasion strategies rely on secreted proteins and small molecules, recent work suggests that lipid virulence factors can also target immune cells. For instance, lipopolysaccharides can trigger inflammasome activation and programmed cell death in macrophages.(16, 17) In the presence of the Pseudomonas quinolone signal (PQS) or under bacterial stress such as lysozyme or antimicrobial treatment, *P. aeruginosa* also produces membrane vesicles formed by blebbing of the outer membrane.(18–22) In addition to activating the inflammasome in macrophages,(23, 24) these vesicles can also directly deliver periplasmic content, including virulence factors, to host cells through membrane fusion.(25, 26) Furthermore, a class of glycolipids called rhamnolipids that are regulated by quorum sensing, has been shown to exert cytotoxicity towards phagocytic cells.(27, 28) The contribution of these various *P. aeruginosa* virulence factors to immune cell death and host immune evasion remain unclear.

In this report, we demonstrate that rhamnolipids predominantly drive *P. aeruginosa* acute virulence against murine macrophages. We show that secreted rhamnolipids can form micelles that exhibit acute cytotoxicity, rupturing the macrophage plasma membrane and damaging intracellular organellar membranes within minutes. We also examine these rhamnolipid micelles’ structural and biochemical properties via transmission electron microscopy and liquid chromatography-mass spectrometry. Furthermore, we demonstrate that while these micelles are particularly toxic to macrophages, they are also capable of damaging a wide range of other cells, including human bronchial epithelial cells, red blood cells, and even Gram-positive bacteria. Finally, we report that rhamnolipid production in various panels of clinical isolates strongly correlates with *P. aeruginosa* virulence.

## Results

### Lipid-rich material secreted by *P. aeruginosa* is toxic to murine macrophages

We previously showed that low-molecular weight secreted material from *P. aeruginosa* is cytotoxic to murine macrophages and that this toxicity is partially dependent on siderophore pyoverdine.(12) Higher-molecular weight materials such as *Pseudomonas* Exotoxin A and protease IV have also been known to kill these cells,(29–32) but a comprehensive evaluation of the relative impact of secreted virulence factors has not yet been performed. To determine which secreted factors contribute to virulence against murine macrophages, RAW264.7 cells were treated with supernatant from bacteria grown in modified M9-casamino acid medium, which was previously used to study the acute virulence of *P. aeruginosa* against the nematode host *Caenorhabditis elegans*.(33) Supernatants from these growth cultures were highly toxic to RAW264.7 cells, causing nearly complete cell death within 5 h **(Fig. 1A)**. Based on our previous studies,(12, 33, 34) a pyoverdine biosynthetic mutant PA14*pvdF* was used to test whether toxicity was mediated by pyoverdine. Loss of pyoverdine biosynthesis by this mutant did not significantly attenuate cell death **(Fig. 1A)**.

**Fig. 1.**
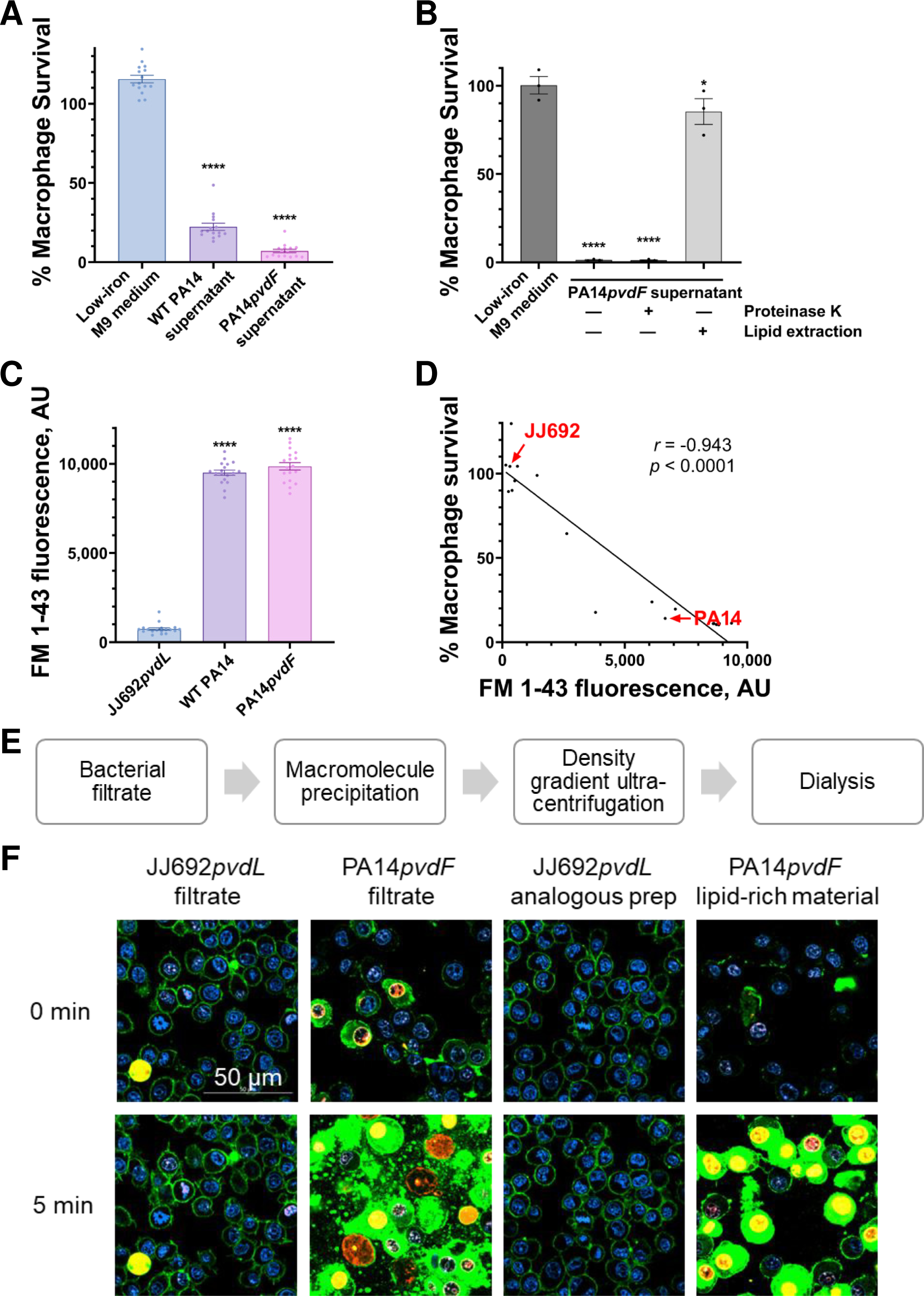
Lipid-rich material in the *P. aeruginosa* supernatant is highly toxic to murine macrophages. **(A)** Murine macrophage (RAW264.7) survival after exposure to supernatants from wild-type PA14 and PA14*pvdF* (grown in low-iron M9 medium). Survival was normalized to saline control. **(B)** Macrophage survival after exposure to PA14*pvdF* supernatants that have been pre-treated with 100 µg/mL Proteinase K (inactivated by 5 mM PMSF) or depleted of lipids by chloroform extraction. Survival was normalized to saline control. **(C)** Quantification of lipids in the supernatant by fluorescence after 20 µg/mL FM 1-43 treatment. **(D)** Correlation between FM 1-43 fluorescence and toxicity towards murine macrophages for supernatants of 19 *P. aeruginosa* clinical and environmental isolates. Representative strains with high lipid content (PA14) and low lipid content (JJ692) are labeled in red. Survival was normalized to media control. **(E)** Schematic of purification pipeline for lipid-rich material. **(F)** Interactions between RAW264.7 cells and bacterial filtrate or purified lipid-rich material from PA14*pvdF* or JJ692*pvdL* in the presence of SYTOX Orange cell-impermeant nucleic acid stain [red]. Secreted bacterial lipids were prelabeled with FM 1-43 [green]. Cells were prelabeled with Hoechst 33342 cell-permeant nucleic acid stain [blue]. Data in **A, B, C** were analyzed via one-way ANOVA. *, *p* < 0.05; **, *p* < 0.01; ***, *p* < 0.001; ****, *p* < 0.0001; ns, not statistically significant.

Material from this mutant was used to further investigate which class(es) of secreted macromolecules (e.g., proteins, lipids) were responsible for cytotoxicity. Spent medium was treated with proteinase K or by using chloroform to extract lipids. Proteinase K treatment had no apparent effect on the toxicity of the spent medium **(Fig. 1B)**, suggesting proteins were not responsible for the cell death. In contrast, lipid extraction dramatically reduced cytotoxicity, indicating that cell death was associated with a lipidaceous virulence factor.

FM 1-43, a probe that remains nonfluorescent in aqueous solution but becomes highly fluorescent when bound to lipid membranes or vesicles,(35–37) was used to assess the presence of lipids in the supernatant. As expected, supernatants from both wild-type PA14 and PA14*pvdF* showed high FM 1-43 fluorescence, indicating the presence of substantial lipid **(Fig. 1C)**. Not all *P. aeruginosa* isolates secreted considerable amounts of lipids; the urinary tract infection isolate JJ692 had nearly 100-fold less FM 1-43 signal (**Fig. 1C**).(38) We surveyed supernatant cytotoxicity and lipid content, as indicated by FM 1-43 fluorescence, in a well-characterized panel of 19 *P. aeruginosa* clinical and environmental isolates.(39) A strong, negative correlation (*r* = - 0.943) was observed in these strains between macrophage survival and the amount of secreted lipids **(Fig. 1D)**. Approximately half of the strains, including JJ692 **(Fig. 1D** – labeled in red), exhibited minimal virulence towards RAW264.7 cells.

To characterize this unknown toxic lipid, a purification pipeline was developed **(Fig. 1E; Fig. S1A)**. First, the bacterial filtrate was mixed with ammonium sulfate. After mixing and centrifugation, low-density floc was observed floating on top of the PA14*pvdF* filtrate **(Fig. S1B).** This material was collected and mixed with an equal concentration of 80% (w/v) Nycodenz and the resulting material was put onto a discontinuous Nycodenz step gradient. Lipid material was collected from the gradient and reconstituted in PBS prior to further use.

Floc was absent from identically-treated material from the negative control strain (JJ692*pvdL*). Instead, the latter strain formed a distinct pellet after centrifugation. The liquid at the top layer of JJ692*pvdL* sample had virtually no FM 1-43 fluorescence and no toxicity. Hence, we performed further experiments with segregated material (floc for PA14*pvdF* and pellet for JJ692*pvdL*). We collected these respective products and assayed them for lipids using FM 1-43. The purified foam material showed high fluorescence, indicating substantial lipid in the floc. Floc was purified by density gradient ultracentrifugation, which resulted in a distinct layer with high lipid content that was only present in the PA14*pvdF* sample **(Fig. S1C**, highlighted in red).

Confocal laser-scanning microscopy was used to observe the interaction between RAW264.7 cells and this FM 1-43-labeled, lipid-rich material or bacterial filtrate. Both PA14*pvdF* filtrate and purified material killed ∼90% cells within 5 min, as indicated by live cell staining with SYTOX Orange (a cell-impermeant nucleic acid stain) **(Fig. 1F; Fig. S1D)**. Analogous material from JJ692*pvdL* had minimal effect on cell viability. Immediately prior to the appearance of bright SYTOX Orange nuclear staining (i.e., cell death), we observed abrupt internalization of FM 1-43. The membrane dye quickly labeled cellular lipids that were soon released from cells, indicating that the purified material caused acute host membrane permeabilization **(Fig. 1F; Fig. S1D; Movie S1 and S2)**.

### Structural characterization of cytotoxic *P. aeruginosa* micelles

We characterized the purified lipid-rich material and macromolecule floc via transmission electron microscopy (TEM). Due to the differing requirements for sample processing for positive and negative staining of transmission electron microscopy, we applied positive staining for floc (solid) **(Fig. 2A-2C)** and negative staining for purified material (aqueous) **(Fig. 2D and E)**. TEM micrographs revealed micellar structures in both samples that were ∼30 nm in diameter **(Fig. 2F)**. No such structures were found in the prep from JJ692*pvdL* **(Fig. 2G and H)**. Micelles were no longer apparent after chloroform extraction of lipids in PA14*pvdF* material **(Fig. 2I)**; lipid depletion also ameliorated toxicity towards macrophages **(Fig. 1B)**. These results suggest an association between these micelles and the cytotoxic behavior of *P. aeruginosa* spent medium and lipid-rich material.

**Fig. 2.**
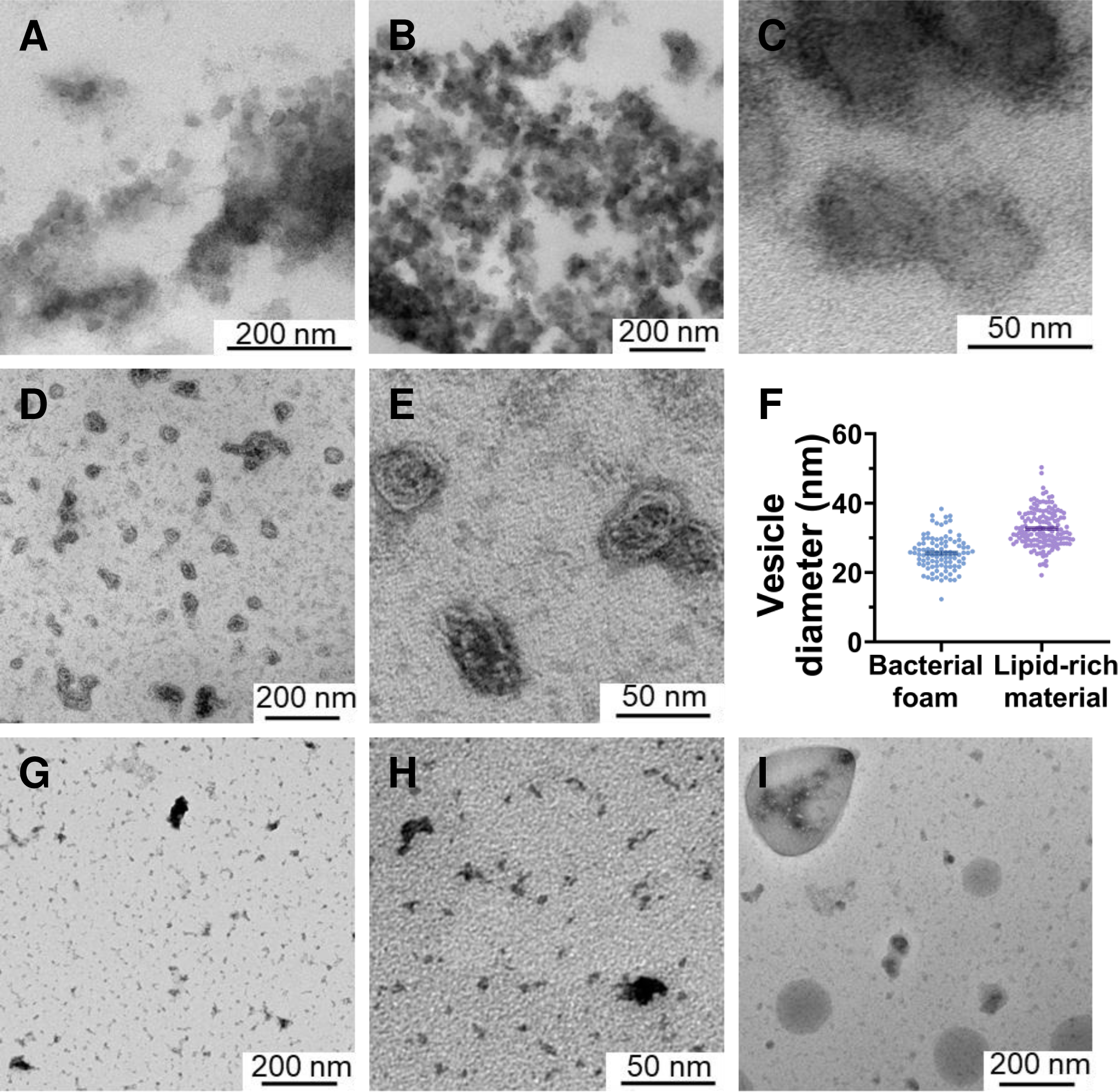
*P. aeruginosa* secretes cytotoxic micelles. **(A, B, C)** Positively-stained macromolecule floc from PA14*pvdF* visualized by transmission electron microscopy (TEM). Digitally zoomed-in view in **C**. **(D, E)** Negatively-stained purified lipid-rich material from PA14*pvdF* visualized by TEM. Digitally zoomed-in view in **E**. **(F)** Average micelle diameter in bacterial floc and lipid-rich material from PA14*pvdF*. **(G, H)** Negatively-stained purified material from JJ692*pvdL.* Digitally zoomed-in view in **H**. **(I)** Negatively-stained purified lipid-rich material from PA14*pvdF* after lipid extraction via chloroform.

### *P. aeruginosa* micelles damage cellular and organellar membranes

We employed TEM to visualize cellular damage during micelle exposure. RAW264.7 cells treated with lipid-deficient material from JJ692*pvdL* showed a clearly identifiable nucleus, intact plasma membrane, and tubular mitochondria **(Fig. 3A-3C)**. In contrast, the majority of cells exposed to purified micelles from PA14*pvdF* exhibited ruptured plasma membranes and severely disrupted mitochondrial membranes, causing the organelles to become engorged **(Fig. 3D-3F)**.

**Fig. 3.**
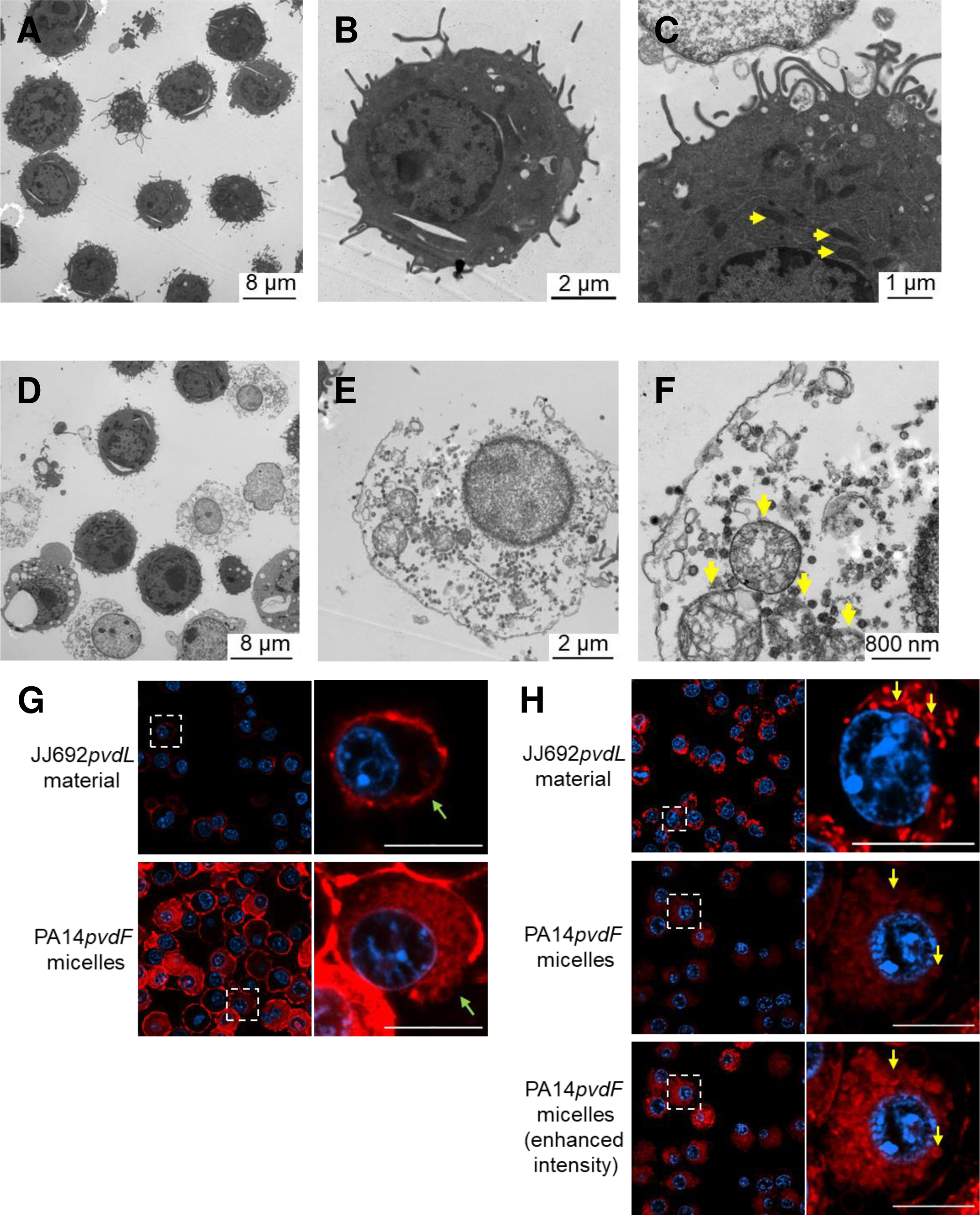
*P. aeruginosa* micelles cause severe damage to the plasma membrane and mitochondrial membrane. **(A-C)** Fixed RAW264.7 cells visualized by transmission electron microscopy (TEM) after exposure to purified sample from JJ692*pvdL*. A representative healthy cell in **B**. Uncompromised mitochondria (yellow arrows) in a healthy cell **(C)**. **(D-F)** Fixed macrophages visualized by TEM after exposure to purified PA14*pvdF*micelles. A representative cell with ruptured plasma membrane in **E**. Detailed view of compromised mitochondria (yellow arrows) in the representative damaged cell **(F)**. **(G)** Visualization of macrophage plasma membrane after 10 min exposure to purified micelles from PA14*pvdF* or material from JJ692*pvdL*. A representative cell (white square) was selected and enhanced for detailed view of the plasma membrane (green arrow). Cells were prelabeled with Hoechst 33342 [blue] and CellMask Deep Red plasma membrane stain [red]. Scale bar = 10 µm. **(H)** Visualization of macrophage mitochondria after 10 min exposure to purified micelles from PA14*pvdF* or material from JJ692*pvdL*. A representative cell (white square) was selected and enhanced for detailed view of individual mitochondria (yellow arrows). Cells were pre-labeled with Hoechst 33342 [blue] and MitoTracker Red CMXRos [red]. Scale bar = 10 µm.

To visualize this phenomenon in real-time, we pre-labeled the plasma membrane and mitochondria using a CellMask deep red plasma membrane stain and MitoTracker Red CMXRos respectively and performed time-course confocal microscopy during micelle exposure. The plasma membrane expanded and ruptured within 10 min **(Fig. 3G; Fig. S2A)**. The compromised plasma membrane was consistent with what was observed in electron micrographs **(Fig. 3E)**. Similarly, mitochondria rapidly fragmented and became engorged upon micelle exposure **(Fig. 3H; Fig. S2B)**. This treatment also significantly reduced MitoTracker Red fluorescence **(Fig. S2C)**, likely due to the loss of mitochondrial membrane potential. Together, these results indicate that *P. aeruginosa* micelles exert their cytotoxicity by damaging cellular and organellar membranes.

### *P. aeruginosa* micelles damage a wide range of host membranes

In addition to murine macrophages, we tested micelle toxicity towards human bronchial epithelial cells (16HBE). PA14*pvdF* filtrate and purified micelles also killed these cells, while lipid-deficient JJ692*pvdL* samples remained largely nontoxic **(Fig. 4A)**. Since these micelles damage plasma membranes, their effects on giant plasma membrane vesicles (GPMVs) derived from 16HBE cells(40) were tested. GPMVs are mainly composed of plasma membrane and limited cytosolic content and thus can be used to study the interactions between *P. aeruginosa* micelles and cellular membranes without triggering cell death pathways that may otherwise cause membrane rupture (e.g., necroptosis).(41, 42)

**Fig. 4.**
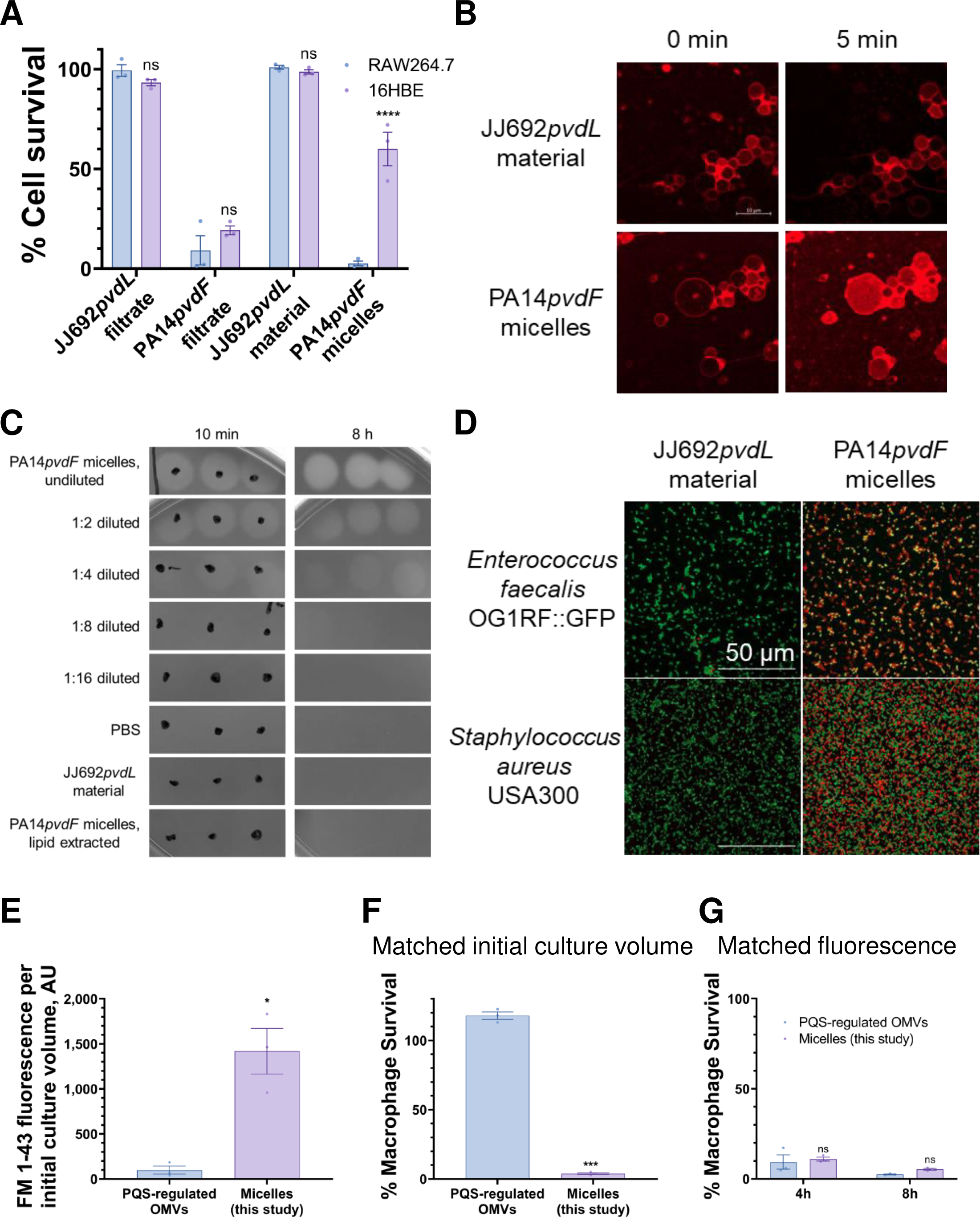
*P. aeruginosa* micelles kill eukaryotic and prokaryotic cells. **(A)** Cytotoxicity of bacterial filtrate and micelles from PA14*pvdF* or material from JJ692*pvdL* against RAW264.7 and human bronchial epithelial cells (16HBE). Data in **A** was analyzed via two-way ANOVA. *, *p* < 0.05; **, *p* < 0.01; ***, *p* < 0.001; ****, *p* < 0.0001; ns, not statistically significant. **(B)** Visualization of giant plasma membrane vesicles (GPMVs) derived from 16HBE cells after 5 min exposure to micelles from PA14*pvdF* or material from JJ692*pvdL*. GPMVs were prelabeled with CellMask Deep Red plasma membrane stain. Scale bar = 10 µm. **(C)** Hemolysis of erythrocytes on sheep blood’s agar after 10 min or 8 h exposure to micelles from PA14*pvdF*, material from JJ692*pvdL*, or PA14*pvdF* sample after lipid extraction via chloroform. **(D)** Visualization of *Enterococcus faecalis* OG1RF::GFP and *Staphylococcus aureus* USA300 after 4 h exposure to micelles from PA14*pvdF* or material from JJ692*pvdL*. **(E-G)** Comparison between pseudomonas quinolone signal (PQS)-regulated outer membrane vesicles (OMVs) and micelles (this study) from wild-type PA14. **(E)** FM 1-43 fluorescence reads in samples after normalized to their initial culture volume respectively. **(F)** Cytotoxicity of samples towards macrophages when matching the fluorescence read. **(G)** Cytotoxicity of samples towards macrophages when matching to the respective initial culture volume. Error bars in **A, E, F, G** represent SEM of 3 biological replicates. Data in **A, F** were analyzed via two-way ANOVA. Data in **E, G** were analyzed via t test. *, *p* < 0.05; **, *p* < 0.01; ***, *p* < 0.001; ****, *p* < 0.0001; ns, not statistically significant.

Purified micelles from PA14*pvdF* ruptured GPMVs within minutes **(Fig. 4B; Fig. S3A)**, suggesting that micelles cause cell death by directly damaging host membranes rather than by triggering host cells to lyse themselves. We further tested micelle toxicity towards erythrocytes using sheep blood agar since these cells lack mitochondria and nuclei. We observed rapid hemolysis within 10 min upon PA14*pvdF* micelle treatment **(Fig. 4C)**. Extracting lipids from this sample or using material from JJ692*pvdL* prevented hemolysis even after 8 h **(Fig. 4C)**. Since *P. aeruginosa* is known to cause hemolysis through the secretion of hemolytic phospholipase C (PlcH), we also tested whether micelles from a *plcH* (PA14*plcH*) mutant could lyse red blood cells. Genetic disruption of *plcH* did not affect the hemolytic properties of purified micelles **(Fig. S3B)**. Moreover, *plcH* mutant still had high FM 1-43 fluorescence of their spent media and corresponding high cytotoxicity toward macrophages **(Fig. S3C)**.

To investigate whether *P. aeruginosa* micelles damaged a wide range of host membranes, including that of noneukaryotic cells, we also measured micelle toxicity towards several microorganisms. The Gram-positive bacteria *Enterococcus faecalis* and *Staphylococcus aureus* are frequently co-isolated with *P. aeruginosa* from biofilms.(43–45) Treatment with PA14*pvdF*-derived micelles caused significant bacterial death in *E. faecalis,* as indicated by the accumulation of propidium iodide staining. Bacteria treated with material from JJ692*pvdL* remained mostly viable **(Fig. 4D)**. Micelles also exhibited bactericidal properties against *S. aureus* **(Fig. 4D),** albeit at a slightly lower rate. However, these micelles showed little activity against *Escherichia coli* and *Candida albicans* **(Fig. S3D)**, likely due to structural differences in the cell walls of Gram-positive bacteria, Gram-negative bacteria, and fungal cells.

### *P. aeruginosa* micelle-mediated damage is distinct from that of PQS-regulated outer membrane vesicles (OMVs)

One major lipidaceous virulence factor of Gram-negative bacteria is outer membrane vesicles (OMVs), which can be produced by *P. aeruginosa* in the presence of the Pseudomonas quinolone signal(20, 22) or under bacterial stress such as lysozyme or antimicrobial treatment.(18, 19, 21) These vesicles can not only activate the inflammasome in macrophages,(23, 24) but also deliver periplasmic content within the community or to host cells through membrane fusion.(25, 26) The OMVs induced by the quorum-sensing molecule 2-heptyl-3-hydroxy-4-quinolone (Pseudomonas quinolone signal, PQS) in BHI medium could package PQS and then traffic this signal molecule within a population, enabling intercellular communication and group behavior.(20, 22) Cytotoxic micelles purified as described were compared to PQS-induced OMVs produced using previously-established, optimized protocols.(46)

Consistent with previous studies, OMVs produced by *P. aeruginosa* under this condition were measurable by FM 1-43 fluorescence (**Fig. 4E**). When normalized to initial culture volume, FM fluorescence of our micelles was 25-30 times higher than that of OMV sample, suggesting that considerably more lipid micelles are produced than OMVs. Micelle cytotoxicity was also considerably higher; >95% of cells were killed, while OMV cytotoxicity was not detected (**Fig. 4F**). If equivalent amounts of fluorescent materials were used, cytotoxic micelles and OMVs both exhibited substantial cytotoxicity **(Fig. 4G)**. These data suggest that, although multiple kinds of bacterial lipids can be damaging to eukaryotic cells if present at sufficient concentration, micelles and OMVs are produced in different concentrations and possibly via different mechanisms.

### Cytotoxic *P. aeruginosa* micelles are composed of rhamnolipids

Previous studies have demonstrated that secreted rhamnolipids from *P. aeruginosa* can induce hemolysis(47, 48), as was observed for the purified micelles (**Fig. 4C**). To ascertain whether these cytotoxic micelles were comprised of rhamnolipids, partially purified micelles were analyzed via liquid chromatography-mass spectrometry (LC-MS) and compared to two commercially-sourced rhamnolipid standards, one enriched in mono-rhamnolipids and the other in di-rhamnolipids **(Fig. 5A)**.

**Fig. 5.**
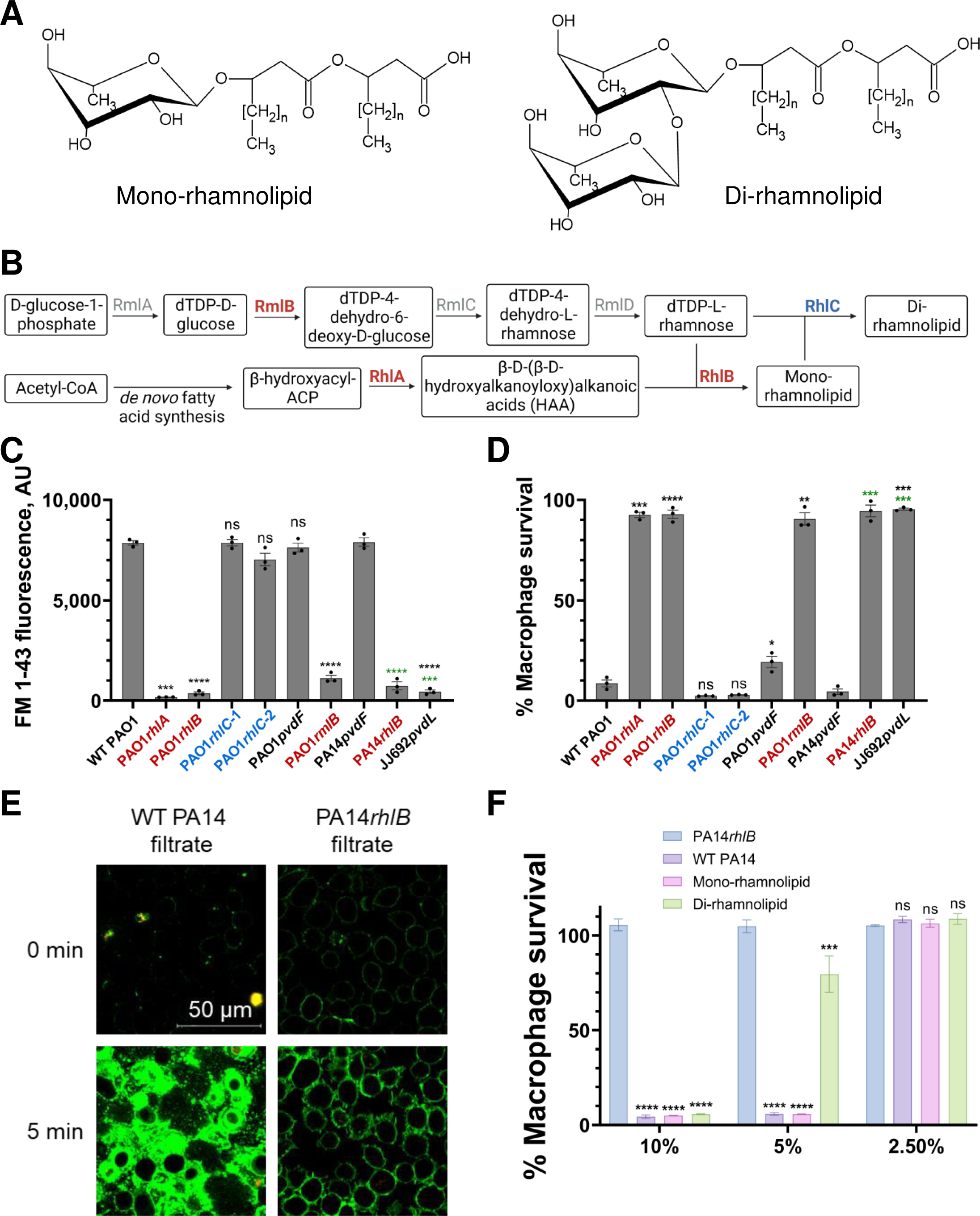
Rhamnolipid biosynthetic mutants do not produce cytotoxic micelles. **(A)** Chemical structures of mono-rhamnolipids and di-rhamnolipids. **(B)** Rhamnolipid biosynthetic pathway in *P. aeruginosa*. **(C)** Lipid content in supernatants from PAO1 and PA14 transposon mutants measured by FM 1-43 fluorescence. **(D)** Cytotoxicity of supernatants from PAO1 and PA14 transposon mutants against RAW264.7 cells. Black labels indicate statistical significance compared to wild-type PAO1. Green labels indicate statistical significance compared to PA14*pvdF*. **(E)** Interactions between RAW264.7 cells and bacterial filtrate from wild-type PA14 or PA14*rhlB* in the presence of SYTOX Orange cell-impermeant nucleic acid stain [red]. Secreted bacterial lipids were prelabeled with FM 1-43 [green]. **(F)** Cytotoxicity of two commercially-sourced rhamnolipid products (enriched in mono-rhamnolipids and di-rhamnolipids respectively) at rhamnolipid concentrations comparable to wild-type PA14 filtrate (standardized by FM 1-43 fluorescence). PA14*rhlB* filtrate was volume-matched to wild-type PA14 filtrate. Error bars in **C, D, F** represent SEM of 3 biological replicates. Data in **C, D** were analyzed via one-way ANOVA. Data in **F** was analyzed via two-way ANOVA. *, *p* < 0.05; **, *p* < 0.01; ***, *p* < 0.001; ****, *p* < 0.0001; ns, not statistically significant.

More than 60% of the compounds detected in the sample were rhamnolipids **(Table 1)**, two of which corresponded to the predominant species found in the standards (Rha-C10-C10 and Rha-Rha-C10-C10). Analysis identified only a limited number of common membrane lipids detected in LC-MS **(Table S1)**, indicating that these micelles were not derived from membranes.

**Table 1.** Cytotoxic *P. aeruginosa* micelles are composed of rhamnolipids.

| Rank | m/z | Mass | Molecular formula | Rhamnolipid components | Overall Vol% | Relative Amount% |
| --- | --- | --- | --- | --- | --- | --- |
| 1 | 531.35 | 532.36 | C<sub>28</sub>H<sub>52</sub>O<sub>9</sub> | Rha-C10-C12/Rha-C12-C10 | 8.98 | 14.52 |
| 2 | 649.38 | 650.39 | C<sub>32</sub>H<sub>58</sub>O<sub>13</sub> | Rha-Rha-C10-C10 | 8.01 | 12.95 |
| 3 | 529.34 | 530.35 | C28H50O9 | Rha-C10-C12:1/Rha-C12:1-C10 | 7.49 | 12.11 |
| 4 | 503.33 | 504.33 | C26H48O9 | Rha-C10-C10 | 6.94 | 11.22 |
| 5 | 677.41 | 678.42 | C34H62O13 | Rha-Rha-C10-C12/Rha-Rha-C12-C10 | 6.87 | 11.11 |
| 6 | 517.34 | 518.35 | C27H50O9 | Rha-C10-C10-CH3 | 6.23 | 10.07 |
| 7 | 475.29 | 476.30 | C24H44O9 | Rha-C8-C10/Rha-C10-C8 | 4.82 | 7.79 |
| 8 | 675.40 | 676.40 | C34H60O13 | Rha-Rha-C10-C12:1/Rha-Rha-C12:1-C10 | 3.22 | 5.21 |
| 9 | 489.31 | 490.31 | C25H46O9 | Rha-C9-C10/Rha-C10-C9 | 2.04 | 3.30 |
| 10 | 663.40 | 664.40 | C33H60O13 | Rha-Rha-C10-C10-CH3/Rha-Rha-C10-C11 | 2.03 | 3.28 |
| 11 | 557.37 | 558.37 | C30H54O9 | Rha-C10-C14:1 | 1.56 | 2.52 |
| 12 | 705.44 | 706.45 | C36H66O13 | Rha-Rha-C12-C12 | 1.36 | 2.20 |
| 13 | 559.39 | 560.39 | C30H56O9 | Rha-C12-C12 | 1.15 | 1.86 |
| 14 | 621.35 | 622.36 | C30H54O13 | Rha-Rha-C8-C10/Rha-Rha-C10-C8 | 1.14 | 1.84 |
| **Overall** |  |  |  |  | 61.84 | 100.00 |

To validate these findings, we measured the lipid content and cytotoxicity of spent media from several rhamnolipid biosynthetic mutants from PAO1 and PA14 harboring transposon insertions in *rhlA*, *rhlB*, *rhlC*, *rmlA, rmlB, rmlC,* or *rmlD* (49–53) (data not shown) **(Fig. 5B)**. Amongst these mutants, only *rmlB*, *rhlA* and *rhlB* failed to produce micelles, or secret lipids in general, based on the low levels of FM 1-43 fluorescence. These strains also displayed significantly attenuated toxicity towards murine macrophages **(Fig. 5C and D)**. Interestingly, spent media from *rhlC* mutants, which are unable to convert mono-rhamnolipids to di-rhamnolipids, exhibited neither decreased lipid content nor lowered cytotoxicity **(Fig. 5C and D)**, suggesting that production of mono-rhamnolipids is sufficient for full toxicity. We also studied the effect of PA14*rhlB* filtrate on RAW264.7 cells via confocal microscopy. Unlike wild-type material, PA14*rhlB* filtrate did not rupture the host membrane **(Fig. 5E; Fig. S4A)**. In addition, we processed PA14*rhlB* filtrate through our micelle purification pipeline **(Fig. 1E)**. We observed that the spent medium did not form apparent floc following ammonium sulfate precipitation **(Fig. S4B)** or the distinct layer with high lipid content after ultracentrifugation **(Fig. S4C)**. The fully purified material from the PA14*rhlB* mutant, prepared identically to micelles from wild-type (WT) PA14 or PA14*pvdF,* did not lyse red blood cells **(Fig. S4D)**, affirming that rhamnolipid production is necessary for the secretion of cytotoxic micelles. At rhamnolipid concentrations comparable to that of purified WT PA14 micelles (less than 1 mg/mL, standardized by FM 1-43 fluorescence), commercially-sourced rhamnolipids induced cell death in RAW264.7 macrophages **(Fig. 5F)**.

To investigate whether proteins were involved in micelle formation or contributed to cytotoxicity, LC-MS proteomics were used to analyze partially purified micelles. Only three proteins from *P. aeruginosa* were detected. The most abundant was protease IV **(Table S2)**, the iron-regulated, secreted protease PrpL.(54) However, spent media from PA14*ΔprpL* exhibited cytotoxicity comparable to that of WT PA14 **(Fig. S5)**.

### Rhamnolipid production correlates with *P. aeruginosa* virulence against murine macrophages

Finally, we investigated the clinical utility of targeting rhamnolipid production during *P. aeruginosa* infection. We first surveyed rhamnolipid micelle production and the toxicity of spent medium from 12 hematological isolates.(55) 11 of these isolates produced detectable amounts of lipid micelles, 7 of which were comparable to PA14 and were toxic to murine macrophages, showing 10% survival or less **(Fig. 6A)**. Overall, we observed a strong, significant negative correlation between *P. aeruginosa* supernatant rhamnolipid content and macrophage survival **(Fig. 6A)**. No correlation was observed between bacterial growth and rhamnolipid production **(Fig. S6A)**.

**Fig. 6.**
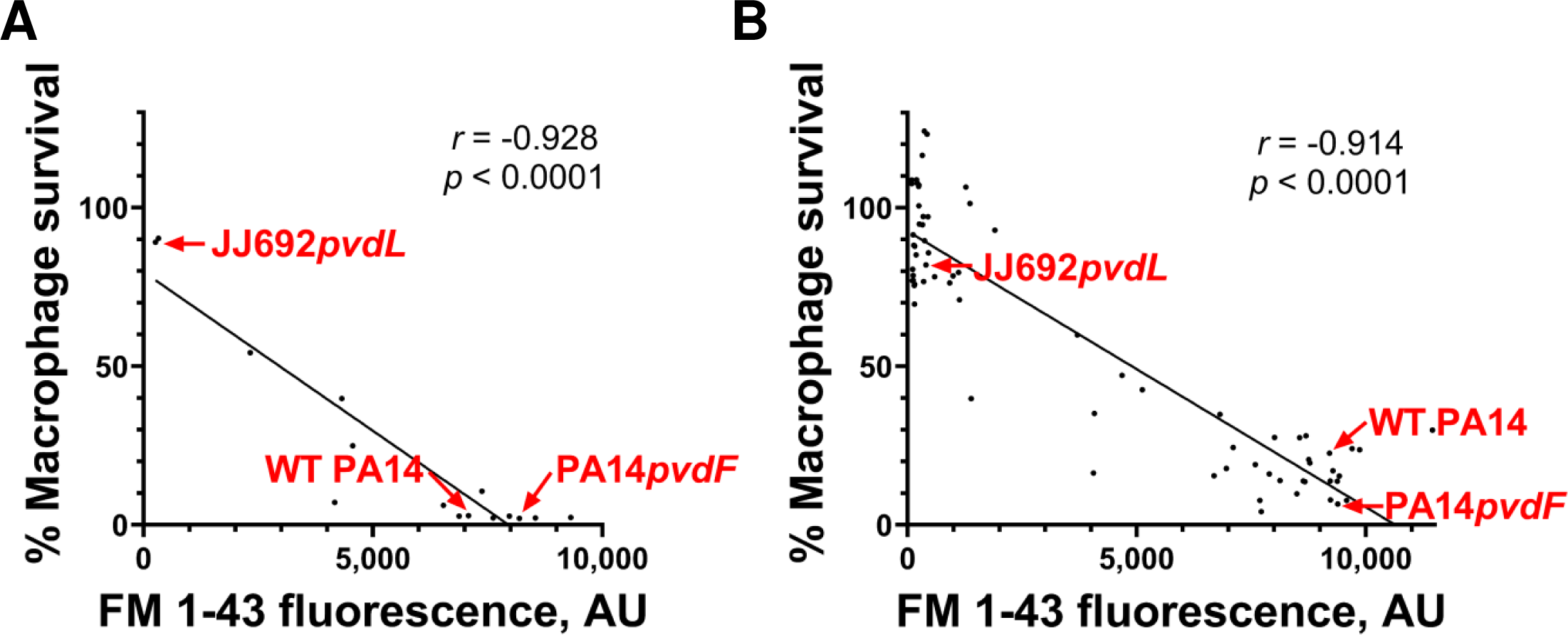
Supernatant rhamnolipid content strongly correlates to cytotoxicity in *P. aeruginosa* clinical isolates. **(A)** Correlation between rhamnolipid micelle production (FM 1-43 fluorescence) and cytotoxicity against murine macrophages for supernatants from 12 hematological isolates. The three control strains are labeled in red. Cell survival was normalized to saline control. **(B)** Correlation between rhamnolipid micelle production (FM 1-43 fluorescence) and cytotoxicity against macrophages for supernatants from 68 clinical isolates from pediatric cystic fibrosis patients. The three control strains are labeled in red. Cell survival was normalized to saline control.

Taking advantage of a larger panel of 68 clinical isolates from pediatric cystic fibrosis patients,(34) this analysis was expanded to a larger group of strains. Nearly half of the isolates in this panel lost rhamnolipid production and also showed little to no toxicity in RAW264.7 cells **(Fig. 6B)**. Similarly, rhamnolipid production strongly correlated to supernatant cytotoxicity, and was independent of bacterial growth **(Fig. S6B)**.

We also performed whole-genome sequencing in selected isolates from these panels and combined these data with pre-existing sequencing data from the Broad Institute(39) for the clinical and environmental isolates we tested in **Fig. 1D**. We compared these strains’ protein sequences for rhamnolipid biosynthetic enzymes (RhlA, RhlB, RhlC, RmlA, RmlB, RmlC, RmlD) and known quorum-sensing regulators (Vfr, LasR, RsaL, LasI, RhlI, RhlR) to the reference strain PAO1. Polymorphisms were found in nearly all of these proteins for both high and low rhamnolipid-producing strains. There was an enrichment in mutations in RhlR mutations (mostly nonsense or frame-shift mutations) amongst isolates that had lost the ability to secrete rhamnolipid micelles **(Table S3)**, suggesting that this quorum-sensing system is a crucial regulator of rhamnolipid micelle production.

## Discussion

In this study, we demonstrated that *P. aeruginosa* secretes rhamnolipids that are highly cytotoxic to murine macrophages and blood cells, rupturing their plasma membranes within minutes. TEM analysis showed that the secreted rhamnolipids formed micelles. Using confocal microscopy and TEM, we observed rhamnolipid-mediated destruction of cellular membranes, including plasma and mitochondrial membranes. We also characterized the structural and biochemical properties of rhamnolipid micelles using TEM and LC-MS. Importantly, rhamnolipid micelles damaged a wide-range of host membranes, including those from murine macrophages, human bronchial epithelial cells, erythrocytes, and Gram-positive bacteria, suggesting that these micelles could also modulate the permeability of the human airway epithelium or influence polymicrobial interactions.(56, 57) Others have also reported that rhamnolipids exhibit broad-spectrum antimicrobial properties against bacterial (e.g., *Klebsiella pneumoniae*, *Listeria monocytogenes*) and fungal (e.g., *Mucor circinelloides*, *Verticillium dahlia*) pathogens,(58–61) even synergizing with conventional antibiotics.(62, 63)

Rhamnolipids, a class of glycolipids produced by *P. aeruginosa*, composed of a rhamnosyl head group and 3-(hydroxyalkanoyloxy)alkanoic acid (HAA) fatty acid tail, have previously been shown to play several roles in virulence. For example, rhamnolipids’ amphiphilic structure allows them to reduce water surface tension, making them a potent biosurfactant.(64) They also facilitate *P. aeruginosa* immune evasion by supporting the development of biofilms,(65) and by inhibiting phagocytosis by macrophages and polymorphonuclear leukocytes, even at sublethal concentrations.(27, 28, 66) Rhamnolipid production has also been associated with the development of ventilator-associated pneumonia.(67)

The rhamnolipids secreted in our study appeared to assemble into micelles. This is consistent with studies that have indicated that rhamnolipids self-assemble into various structures, including micelles, vesicles, lamellar structures, and even mesophases, depending on factors such as concentration, pH, temperature, presence of additives, and sample heterogeneity (congeners).(68, 69) However, the micelles we have characterized here are distinctly smaller (∼30 nm in diameter) than those previously characterized (∼100-1000 nm).(70) This discrepancy may have been due to the different media used to grow the pathogen, the different methods for harvesting, and processing of the secreted products. Despite the difference in size, both groups of micelles have been shown to target and kill *S. aureus*.

Previously, rhamnolipid micelles have been shown to contain small *P. aeruginosa* metabolites.(70) However LC-MS analysis of our partially purified rhamnolipid material for small molecule and proteomic content suggest that these smaller micelles didn’t contain any cargo. Instead, toxicity appears to be a consequence of the lipid itself rather than any encapsulated cargo, though the precise mechanism of this phenomenon remains unclear. It will be important to further elucidate how rhamnolipids interact with host membranes, such as the role of surface glycoproteins or membrane domains (i.e., distribution of cholesterol and sphingolipids) and the molecular basis of membrane rupture. These mechanistic studies could provide some insight into how rhamnolipids relate to *P. aeruginosa* virulence.

Available evidence indicates that the connection of rhamnolipids to virulence appears complex. Since rhamnolipids have been detected in the sputum of cystic fibrosis patients,(71) several studies have reported rhamnolipid production by *P. aeruginosa* cystic fibrosis isolates.(57, 72) However, while most *P. aeruginosa* isolates produce rhamnolipids during the acute, early stages of infection, many gradually lose this ability.(72) Our observations from a panel of multidrug-resistant isolates from pediatric cystic fibrosis patients showed the same pattern: nearly half of the isolates in this panel failed to produce rhamnolipids **(Fig. 6B)**, likely due to mutations in RhlR **(Table S3)**. This is consistent with a well-known phenomenon wherein *P. aeruginosa* undergoes a transition from an acute-to-chronic pattern, often associated with mutations in key quorum-sensing regulators such as LasR or RhlR.(72) This phenomenon has been well-documented in cystic fibrosis patients, where *P. aeruginosa* frequently establishes chronic infections that can persist for decades.(73, 74)

Interestingly, nearly all strains in a different panel of isolates from hematological infections exhibited some level of rhamnolipid production **(Fig. 6A)**. These infections typify acute virulence patterns, as the pathogen requires the secretion of various materials, such as toxins, proteases, elastases, etc. to colonize primary infection sites and to traverse into the bloodstream. Our surveys of hematological and cystic fibrosis isolates, where macrophage death was strongly correlated to rhamnolipid production, indicate that this material may play a crucial role during certain infections.

This result, combined with the previously described virulence roles for rhamnolipids, may indicate that targeting rhamnolipid production could have a beneficial effect, particularly during acute infection. Targeting virulence determinants during infection is a strategy that has increasingly received attention as a promising alternative or supplement to conventional antimicrobials. This has been particularly true in response to the emergence of multi- or even pan-drug resistant strains, which has made treating pseudomonal infections increasingly challenging.

## Materials and Methods

### Bacterial strains and growth conditions

*P. aeruginosa* strain PA14 (wild-type), pyoverdine biosynthetic mutant PA14*pvdF*, and mutant of hemolytic phospholipase C (PA14*plcH*) as well as rhamnolipid biosynthetic mutants (PA14*rhlB*) were all obtained from the UCBPP-PA14 transposon mutant library.(75) The mutant with protease IV deletion PA14*ΔprpL* was obtained from Dr. Frederick Ausubel. *P. aeruginosa* strain PAO1 (wild-type), pyoverdine biosynthetic mutant PAO1*pvdF*, and its rhamnolipid biosynthetic mutants (*rhlA*, *rhlB*, *rhlC*, *rmlA, rmlB, rmlC, and rmlD*) were all obtained from PAO1 two-allele transposon mutant library.(76, 77) JJ692*pvdL* was constructed using the pMAR2xT7 vector containing the mariner transposon as previously described.(75)19 *P. aeruginosa* clinical and environmental isolates in Fig. 1 were from Dr. Frederick Ausubel.(39) 12 *P. aeruginosa* hematological isolates patients were obtained from the OHSU Clinical Microbiology lab after being isolated from BSIs in OHSU HCT/HM patients, provided by Dr. Morgan Hakki.(55) 68 Deidentified *P. aeruginosa* isolates from pediatric cystic fibrosis patients were provided by Dr. Carolyn Cannon.(34)

For all experiments, an overnight culture of *P. aeruginosa* was first grown in LB medium for 12-14 hours. The culture was diluted 1:100 into low-iron M9 medium (1% 5× M9 salts [Difco], 1.3% low-iron Casamino Acids [Difco]) supplemented with 1 mM MgSO_4_ and 1 mM CaCl_2_, and incubated for 16-20 hours (37 °C, 225 rpm). Bacterial growth (absorbance at 600 nm) and FM 1-43 (Invitrogen) fluorescence were measured using a Cytation5 multimode plate reader (BioTek).

To test micelle toxicity towards microorganisms, *Candida albicans* fRS26::GFP, *Enterococcus faecalis* OG1RF::GFP, *Escherichia coli* OP50::GFP, and *Staphylococcus aureus* USA300 were used.(78) fRS26::GFP and OG1RF::GFP were grown in Brain Heart Infusion (BHI) broth for 12-14 hours (37 °C, 225 rpm). OP50::GFP and USA300 were grown in LB medium for 12-14 hours (37 °C, 225 rpm).

### Rhamnolipid micelle purification

For the preparation of bacterial supernatant, the M9 culture was centrifuged at 13,300 rpm for 15 min. Antibiotics, including Amikacin, Carbenicillin, and Tobramycin (final concentration was 100 µg/mL each), were added into the collected supernatant.

Bacterial filtrate was applied for rhamnolipid micelle purification. Similarly, to prepare bacterial filtrate, the overnight M9 culture was first centrifuged at 10,000 rpm for 40 min, later filtered through 0.2 µm PES membrane (Thermo Scientific), and supplemented with antibiotics. The macromolecules inside the filtrate would form a layer of floc floating on the top when adding ammonium sulfate to 75%. This material was collected and mixed with 80% Nycodenz (w/v) to make 40% Nycodenz solution. Nycodenz gradients were then layered into an ultracentrifuge tube at concentrations of 40% (the layer with bacterial material), 20%, 10%, and 0 (PBS only). Gradients were ultracentrifuged at 41,000 rpm for 4 h at 4 °C. The harvested lipid-rich material went through 2h dialysis within 2K molecular-weight cut off cassettes (Thermo Scientific) to be reconstituted into phosphate-buffered saline (PBS). The final product was stored at −80 °C.

For Proteinase K treatment, PA14*pvdF* supernatant was incubated with Proteinase K (Promega) (working concentration: 100 µg/mL) at 37 °C for 24 h. The reaction was later stopped by adding PMSF (working concentration: 5 mM) and incubating at room temperature for 1 h. For lipid extraction, PA14*pvdF* supernatant or rhamnolipid micelles were 1:1 (volume to volume) mixed with chloroform with vortex and later centrifuged at 13,300 rpm for 15 min. The top layer was collected as lipid-extracted material.

### PQS-regulated OMV purification

Pseudomonas quinolone signal (PQS)-regulated outer membrane vesicles (OMVs) were generated from wild-type PA14 as described previously.(46) Briefly, PA14 was grown in BHI broth for 12 hours (37 °C, 250 rpm). The culture was centrifuged at 15,000 ×g at 4°C for 15 minutes to pellet the cells, and the supernatant was filtered through a 0.45 µm syringe filter. The supernatant was centrifuged at 200,000 ×g at 4 °C for 1.5 hours to pellet out the OMVs. The pellet was resuspended in 500 µL of MV Buffer (50mM Tris, 5mM NaCl, 1mM MgSO4, pH 7.4). Those rhamnolipid micelles used for comparison, which were labeled as ‘Micelles (this study)’ were also from wild-type PA14.

### Cell culture

Murine macrophages (RAW264.7) were maintained at 37 °C in 5% CO_2_ in RPMI-1640 medium containing 10% bovine calf serum and 1% Penicillin-Streptomycin (P/S). Human bronchial epidermal cells (16HBE), which have been immortalized by SV40 large T-antigen, were cultured at 37°C in 5% CO_2_ in MEM medium supplemented with 10% fetal bovine serum, 1% non-essential amino acids (NEAA) and 1% P/S.

### Cell viability assay

Viability of each cell was quantified using alamarBlue HS Cell Viability Reagent (Invitrogen). Cells were seeded in a 24-well plate (about 1 million/well) overnight, and used when reaching 80% confluence. Before adding sample to cells, the culture medium was replaced with serum-free medium (RPMI-1640 medium containing 1% P/S, or MEM medium with 1% NEAA and 1% P/S respectively). In most experiments, tested samples were first diluted in PBS to designed concentrations, 150 µL of which was added into 350 µL serum-free medium in each well. The 24-well plate was incubated at 37°C in 5% CO_2_ for 4 h. And then 50 µL alamarBlue reagent was added into each well. After incubation for another 1 h, the fluorescence at 590 nm was measured and normalized to the well with 150 µL PBS. The cytotoxicity data of 19 common isolates were normalized to M9 medium.

For comparison with PQS-regulated OMVs, when matching FM 1-43 fluorescence, 250 µL samples was added into 250 µL serum-free medium in each well. The 24-well plate was incubated at 37°C in 5% CO_2_ for 3 h or 7 h. The cytotoxicity data were normalized to the well with 250 µL MV buffer. When matching initial culture volume, 25 µL rhamnolipid micelle sample or 42 µL OMV sample was added into serum-free medium in each well (final volume: 500 µL). The 24-well plate was incubated at 37°C in 5% CO_2_ for only 1 h. The cytotoxicity data were normalized to the well with 42 µL MV buffer.

### Fluorescence imaging

The fluorescence images of cells were taken via Zeiss LSM800 Airyscan fluorescence microscopy. RAW264.7 cells were seeded in an 8-well plate (about 0.75 million/well) overnight, and used when reaching 90% confluence. The cell medium for imaging was similar to the one for viability assay, 70% serum-free medium and 30% test sample at designed concentrations.

The working concentrations of each dye for cells: Hoechst 33342 (Thermo Scientific), 40 µM in serum-free medium when imaging; SYTOX Orange (Invitrogen), 10 µM in serum-free medium when imaging; CellMask deep red plasma membrane stain (Invitrogen), 10 µg/mL in serum-free medium (rinsed before imaging); MitoTracker Red CMXRos (Invitrogen), 1 µM in serum-free medium medium (for cells, rinsed before imaging). The working concentrations of FM 1-43 to pre-stain rhamnolipid micelle samples (including bacterial supernatant/ filtrate, purified rhamnolipid micelles or JJ692*pvdL* material etc.) was 20 µg/mL.

The actual color of SYTOX Orange is orange (Ex: 547 nm, Em: 570 nm), while that of FM 1-43 is orange as well (Ex: 473 nm, Em: 579 nm). Here we assigned SYTOX Orange as red and FM 1-43 as green to avoid confusion in images.

### Transmission Electron Microscopy (TEM)

For PA14*pvdF* micelles and JJ692*pvdL* material, as well as lipid-extracted micelles, freshly made samples using the purification pipeline above were negatively stained with 3% uranyl acetate (Electron Microscopy Sciences [EMS], Cat# 22400). For TEM of macrophages, cells were incubated with PA14*pvdF* micelles and JJ692*pvdL* material for one minute, fixed with Karnovsky’s Fixative (EMS, Cat# 15732-10) overnight and pelleted for TEM analysis. Pelleted samples were then post-fixed for one hour in 1% osmium tetroxide (EMS, Cat# 19100), dehydrated in a graded series of ethanol, embedded in Embed812 epoxy resin (EMS, Cat# 14120) and heat polymerized overnight at 70 °C. Samples were then sectioned at 100 nm thickness using a Leica EM UC7 ultramicrotome. Sections were positively stained with saturated methanolic uranyl acetate and Reynold’s lead citrate (EMS, Cat# 22410). Both negatively and positively stained samples were imaged using a JEOL JEM-1230 TEM operating with 80 kV of accelerating voltage and equipped with an AMT NanoSprint15 mKII sCMOS camera.

### Toxicity towards microorganism

Toxicity of rhamnolipid micelles towards microorganisms was visualized using a fluorescent microscope (Zeiss Axio Imager M2). 25 µL rhamnolipid micelles were added into 75 µL 12-14 h microorganism culture in respective media. The mixture was incubated at 37°C with 225 rpm shaking for 4 h and then centrifuged at 13,300 rpm for 5 min. The pellet was resuspended in 25 µL PBS with 2 µg/mL propidium iodide for OG1RF::GFP and USA300 or 25 µL for OP50::GFP. The pellet of USA 300 was resuspended in 50 µL PBS with 40 µM acridine orange and 2 µg/mL propidium iodide. Micrographs of microorganisms were taken via Zeiss Axio Imager M2 fluorescence microscopy. Three biological replicates were performed.

### GPMV formation

Giant plasma membrane vesicles (GPMV) were generated from 16HBE cells as described previously.(40) Briefly, these cells were first gently rinsed with vesiculation buffer (150 mM NaCl, 2 mM CaCl_2_, and 20 mM HEPES in water, pH 7.4) twice, incubated with active vesiculation buffer (1.9 mM DTT, 27.6 mM formaldehyde (HCHO), in vesiculation buffer) for 4 h and followed by centrifugation at 500 ×g for 5 min to remove cellular debris.

To concentrate GPMVs, these vesicles were stained with 5 µg/mL CellMask deep red plasma membrane stain (Invitrogen) and later centrifuged at 13,300 rpm for 15 min. The supernatant was aspirated while the pellet was resuspended in PBS and ready for fluorescence imaging.

### Blood Agar Culture

Blood agar (TSA with sheep blood) medium (Thermo Scientific) was utilized here. Aqueous sample like micelles was dropped to the surface (8 µL each droplet). PBS here was used for dilution and also dropped on the surface. After droplets dried up, the plate was transferred to 37 °C and incubated for 8 h.

### Proteomics study and rhamnolipid study

Partially purified rhamnolipid micelles (floc) were prepared as described above and utilized for liquid chromatography-mass spectrometry (LCMS) analysis. All LC-MS analysis was carried out on an Agilent 6545XT qToF mass spectrometer that was interfaced to an Agilent 1290 Infinity ii chromatography system through a Jet Stream Electrospray Ionization (ESI) source.

For proteomic analysis samples were diluted to a final concentration of ∼ 1 µg/µL in 100 mM ammonium bicarbonate buffer (pH ∼ 7.8). Samples were reduced with tris(2-carboxyethyl)phosphine (TCEP) at 55 °C for 1 h and then alkylated with iodoacetamide (IAA) at room temperature for 1 h, protected from light. Samples were then digested with Trypsin/LysC (Promega, Madison, WI) overnight for ∼ 18 h and subjected to LC-MS/MS analysis. The LC injection volume was 20 µL corresponding to ∼ 20 µg of total digest loaded on column. The LC separations were carried out on a HALO 160 Å ES-C18, 2 µm, 2.1 x 150 mm column operated at 35 °C and a flow rate of 0.4 mL/min. Mobile phase A was 0.1% formic acid in water and mobile phase B was 0.1% formic acid in acetonitrile. The gradient was run from 2% B to 95% B over 86 min as follows: initial conditions 2% B held at 2% B from 0 - 2.5 min, 2.5 - 5 min 8% B, 5 – 33 min 15% B, 33 - 73 min 35% B, 73 - 79 min 65% B, 79 - 82 min 95% B, held at 95% B until 86min. At the end of the run the column was re-equilibrated to initial conditions (2% B) for 4 min. MS Data was collected in the positive ionization mode using AutoMS2. MS data was collected over a range of 300 – 1500 m/z at a scan rate of 8 spectra/sec. MS/MS data was collected over a range of 100 – 1700 m/z at a scan rate of 4 spectra/sec and an isolation width of 4 amu. Collision energy was selected by the MassHunter acquisition software based on z and m/z values. Data analysis and protein database searching was carried out using MASCOT v. 2.7 (Matrix Science, London, UK). Searches were run against the UniRef100 database restricted by taxonomy to Bacteria (Eubacteria).

For rhamnolipid analysis samples were diluted to a final concentration of ∼ 0.5 µg/µL in 10 mM ammonium formate buffer (pH ∼ 3.5). The LC injection volume was 2 µL corresponding to ∼ 1 µg of total sample on column. The LC separations were carried out on an Agilent Eclipse Plus C18 RRHD, 1.8 µm, 2.1 x 50 mm column operated at 40 °C and a flow rate of 0.4 mL/min. Mobile phase A was 10 mM ammonium formate in water (pH ∼ 3.5) and mobile phase B was methanol. The gradient was run from 25% B to 95% B over 25 min as follows: initial conditions 25% B, 0 - 20 min 95% B, 20 - 25 min hold 95% B. At the end of the run the column was re-equilibrated to initial conditions (25% B) for 3 min. MS Data was collected in the negative ionization mode. MS data was collected over a range of 100 – 1100 m/z at a scan rate of 6 spectra/sec. MS/MS data was collected over a range of 75 – 1100 m/z at a scan rate of 2 spectra/sec and an isolation width of 4 amu. Collision energies of 10, 20, and 30 were used for MS/MS data acquisition. Data analysis and database searching was carried out using MassHunter Qualitative Analysis v.10. Initial rhamnolipids were identified by searching the Agilent METLIN PCD Lipids Database v.8. Additional rhamnolipids were identified through manual data inspection.

## Acknowledgements

We extend our gratitude to Dr. Frederick Ausubel for providing *P. aeruginosa* clinical and environmental isolates, Dr. Morgan Hakki for providing *P. aeruginosa* hematological isolates, and Dr. Carolyn Cannon for providing deidentified *P. aeruginosa* isolates from pediatric cystic fibrosis patients. We extend our gratitude to Emily Zhou for her help in surveying 68 clinical isolates. Fig. 5B and Fig. S1A were created with BioRender.com.

This study was supported by the National Institutes of Health (R35GM129294 to NVK), Cystic Fibrosis Foundation (KIRIEN20I0 to NVK, XU23H0 to QX, and KANG19H0, KANG22H0 to DK), and American Heart Association (903591 to DK). The funders had no role in study design, data collection and analysis, decision to publish, or preparation of the manuscript.

## Author contributions

Conceptualization, Q.X., D.K., and N.V.K.; Methodology, Q.X., D.K., and N.V.K.; Investigation, Q.X., D.K., M.D.M., C.L.P., C.G., and J.W.S.; Writing – Original Draft, Q.X. and D.K.; Writing – Review & Editing, Q.X., D.K., M.D.M., C.L.P., C.G., J.W.S., and N.V.K.; Funding Acquisition, Q.X., D.K., and N.V.K.; Resources, C.G. and J.W.S.; Supervision, N.V.K..

## References

1. Gaynes R, Edwards JR, National Nosocomial Infections Surveillance S. 2005. Overview of nosocomial infections caused by gram-negative bacilli. Clinical Infectious Diseases: An Official Publication of the Infectious Diseases Society of America 41:848–854.

2. Spagnolo AM, Sartini M, Cristina ML. 2021. Pseudomonas aeruginosa in the healthcare facility setting. Reviews and Research in Medical Microbiology 32:169–175.

3. Shafran N, Shafran I, Ben-Zvi H, Sofer S, Sheena L, Krause I, Shlomai A, Goldberg E, Sklan EH. 2021. Secondary bacterial infection in COVID-19 patients is a stronger predictor for death compared to influenza patients. Scientific Reports 11:12703.

4. Moradali MF, Ghods S, Rehm BHA. 2017. Pseudomonas aeruginosa Lifestyle: A Paradigm for Adaptation, Survival, and Persistence. Frontiers in cellular and infection microbiology 7.

5. Qin S, Xiao W, Zhou C, Pu Q, Deng X, Lan L, Liang H, Song X, Wu M. 2022. Pseudomonas aeruginosa: pathogenesis, virulence factors, antibiotic resistance, interaction with host, technology advances and emerging therapeutics. Signal Transduction and Targeted Therapy 7.

6. Casilag F, Lorenz A, Krueger J, Klawonn F, Weiss S, Häussler S. 2016. The LasB Elastase of Pseudomonas aeruginosa Acts in Concert with Alkaline Protease AprA To Prevent Flagellin-Mediated Immune Recognition. Infection and Immunity 84:162–171.

7. Jurado-Martín I, Sainz-Mejías M, Mcclean S. 2021. Pseudomonas aeruginosa: An Audacious Pathogen with an Adaptable Arsenal of Virulence Factors. International Journal of Molecular Sciences 22:3128.

8. Bainbridge T, Fick RB. 1989. Functional importance of cystic fibrosis immunoglobulin G fragments generated by Pseudomonas aeruginosa elastase. The Journal of laboratory and clinical medicine 114:728–733.

9. Prasad ASB, Shruptha P, Prabhu V, Srujan C, Nayak UY, Anuradha CKR, Ramachandra L, Keerthana P, Joshi MB, Murali TS, Satyamoorthy K. 2020. Pseudomonas aeruginosa virulence proteins pseudolysin and protease IV impede cutaneous wound healing. Laboratory Investigation 100:1532–1550.

10. Hauser AR. 2009. The type III secretion system of Pseudomonas aeruginosa: infection by injection. Nature Reviews Microbiology 7:654–665.

11. Lovewell RR, Patankar YR, Berwin B. 2014. Mechanisms of phagocytosis and host clearance of Pseudomonas aeruginosa. American journal of physiology 306:L591–L603.

12. Kang D, Kirienko NV. 2020. An In Vitro Cell Culture Model for Pyoverdine-Mediated Virulence. Pathogens 10:9.

13. Azghani AO, Idell S, Bains M, Hancock REW. 2002. Pseudomonas aeruginosa outer membrane protein F is an adhesin in bacterial binding to lung epithelial cells in culture. Microbial pathogenesis 33:109–114.

14. Paulsson M, Su Y-C, Ringwood T, Uddén F, Riesbeck K. 2019. Pseudomonas aeruginosa uses multiple receptors for adherence to laminin during infection of the respiratory tract and skin wounds. Scientific Reports 9.

15. Stoner SN, Baty JJ, Scoffield JA. 2022. Pseudomonas aeruginosa polysaccharide Psl supports airway microbial community development. The ISME Journal 16:1730–1739.

16. Sutterwala FS, Mijares LA, Li L, Ogura Y, Kazmierczak BI, Flavell RA. 2007. Immune recognition of *Pseudomonas aeruginosa* mediated by the IPAF/NLRC4 inflammasome. The Journal of Experimental Medicine 204:3235–3245.

17. Pier G. 2007. Pseudomonas aeruginosa lipopolysaccharide: A major virulence factor, initiator of inflammation and target for effective immunity. International Journal of Medical Microbiology 297:277–295.

18. Kadurugamuwa JL, Beveridge TJ. 1995. Virulence factors are released from Pseudomonas aeruginosa in association with membrane vesicles during normal growth and exposure to gentamicin: a novel mechanism of enzyme secretion. Journal of Bacteriology 177:3998–4008.

19. Kadurugamuwa JL, Beveridge TJ. 1996. Bacteriolytic effect of membrane vesicles from Pseudomonas aeruginosa on other bacteria including pathogens: conceptually new antibiotics. Journal of Bacteriology 178:2767–2774.

20. Mashburn LM, Whiteley M. 2005. Membrane vesicles traffic signals and facilitate group activities in a prokaryote. Nature 437:422–425.

21. Metruccio MME, Evans DJ, Gabriel MM, Kadurugamuwa JL, Fleiszig SMJ. 2016. Pseudomonas aeruginosa Outer Membrane Vesicles Triggered by Human Mucosal Fluid and Lysozyme Can Prime Host Tissue Surfaces for Bacterial Adhesion. Frontiers in Microbiology 7:871.

22. Schertzer JW, Whiteley M. 2012. A Bilayer-Couple Model of Bacterial Outer Membrane Vesicle Biogenesis. mBio 3:e00297-11-e00297.

23. Cecil JD, O’Brien-Simpson NM, Lenzo JC, Holden JA, Singleton W, Perez-Gonzalez A, Mansell A, Reynolds EC. 2017. Outer Membrane Vesicles Prime and Activate Macrophage Inflammasomes and Cytokine Secretion In Vitro and In Vivo. Frontiers in immunology 8.

24. Finethy R, Luoma S, Orench-Rivera N, Feeley EM, Haldar AK, Yamamoto M, Kanneganti T-D, Kuehn MJ, Coers J, Weiss DS, Rubin EJ. 2017. Inflammasome Activation by Bacterial Outer Membrane Vesicles Requires Guanylate Binding Proteins. mBio 8.

25. Berleman J, Auer M. 2013. The role of bacterial outer membrane vesicles for intra- and interspecies delivery. Environmental Microbiology 15:347–354.

26. Schwechheimer C, Kuehn MJ. 2015. Outer-membrane vesicles from Gram-negative bacteria: biogenesis and functions. Nature Reviews Microbiology 13:605–619.

27. McClure CD, Schiller NL. 1992. Effects of Pseudomonas aeruginosa rhamnolipids on human monocyte-derived macrophages. Journal of Leukocyte Biology 51:97–102.

28. McClure CD, Schiller NL. 1996. Inhibition of Macrophage Phagocytosis by Pseudomonas aeruginosa Rhamnolipids In Vitro and In Vivo. Current Microbiology 33:109–117.

29. Pollack M. 1983. The Role of Exotoxin A in Pseudomonas Disease and Immunity. Clinical infectious diseases 5:S979–S984.

30. Michalska M, Wolf P. 2015. Pseudomonas Exotoxin A: optimized by evolution for effective killing. Frontiers in Microbiology 6:963.

31. Engel LS, Hill JM, Moreau JM, Green LC, Hobden JA, O’Callaghan RJ. 1998. Pseudomonas aeruginosa protease IV produces corneal damage and contributes to bacterial virulence. Investigative ophthalmology & visual science 39:662–665.

32. Engel LS, Hill JM, Caballero AR, Green LC, O’Callaghan RJ. 1998. Protease IV, a Unique Extracellular Protease and Virulence Factor from Pseudomonas aeruginosa. Journal of Biological Chemistry 273:16792–16797.

33. Kang D, Kirienko DR, Webster P, Fisher AL, Kirienko NV. 2018. Pyoverdine, a siderophore from Pseudomonas aeruginosa, translocates into C. elegans, removes iron, and activates a distinct host response. Virulence 9:804–817.

34. Kang D, Revtovich AV, Chen Q, Shah KN, Cannon CL, Kirienko NV. 2019. Pyoverdine-Dependent Virulence of Pseudomonas aeruginosa Isolates From Cystic Fibrosis Patients. Frontiers in Microbiology 10.

35. Niles W. 1999. Changes in Fluorescence Spectra of the Dye FM 1-43 in Different Environments. Microscopy Today 7:45–45.

36. Harata N, Ryan TA, Smith SJ, Buchanan J, Tsien RW. 2001. Visualizing recycling synaptic vesicles in hippocampal neurons by FM 1-43 photoconversion. Proceedings of the National Academy of Sciences 98:12748–12753.

37. Richards DA, Bai J, Chapman ER. 2005. Two modes of exocytosis at hippocampal synapses revealed by rate of FM1-43 efflux from individual vesicles. The Journal of Cell Biology 168:929–939.

38. Wolfgang MC, Kulasekara BR, Liang X, Boyd D, Wu K, Yang Q, Miyada CG, Lory S. 2003. Conservation of genome content and virulence determinants among clinical and environmental isolates of Pseudomonas aeruginosa. Proceedings of the National Academy of Sciences 100:8484–8489.

39. Lee DG, Urbach JM, Wu G, Liberati NT, Feinbaum RL, Miyata S, Diggins LT, He J, Saucier M, Déziel E, Friedman L, Li L, Grills G, Montgomery K, Kucherlapati R, Rahme LG, Ausubel FM. 2006. Genomic analysis reveals that Pseudomonas aeruginosa virulence is combinatorial. Genome Biology 7:R90.

40. Gerstle Z, Desai R, Veatch SL. 2018. Giant Plasma Membrane Vesicles: An Experimental Tool for Probing the Effects of Drugs and Other Conditions on Membrane Domain Stability. Methods in Enzymology 603:129–150.

41. Zhang Y, Chen X, Gueydan C, Han J. 2018. Plasma membrane changes during programmed cell deaths. Cell Research 28:9–21.

42. Dhuriya YK, Sharma D. 2018. Necroptosis: a regulated inflammatory mode of cell death. Journal of Neuroinflammation 15.

43. Gjødsbøl K, Christensen JJ, Karlsmark T, Jørgensen B, Klein BM, Krogfelt KA. 2006. Multiple bacterial species reside in chronic wounds: a longitudinal study. International Wound Journal 3:225–231.

44. Korgaonkar A, Trivedi U, Rumbaugh KP, Whiteley M. 2013. Community surveillance enhances Pseudomonas aeruginosa virulence during polymicrobial infection. Proceedings of the National Academy of Sciences 110:1059–1064.

45. Deleon S, Clinton A, Fowler H, Everett J, Horswill AR, Rumbaugh KP. 2014. Synergistic Interactions of Pseudomonas aeruginosa and Staphylococcus aureus in an In Vitro Wound Model. Infection and Immunity 82:4718–4728.

46. Cooke AC, Florez C, Dunshee EB, Lieber AD, Terry ML, Light CJ, Schertzer JW. 2020. Pseudomonas Quinolone Signal-Induced Outer Membrane Vesicles Enhance Biofilm Dispersion in Pseudomonas aeruginosa. mSphere 5:10.1128/msphere.01109-20.

47. Johnson MK, Boese-Marrazzo D. 1980. Production and properties of heat-stable extracellular hemolysin from Pseudomonas aeruginosa. Infection and Immunity 29:1028–1033.

48. Fujita K, Akino T, Yoshioka H. 1988. Characteristics of heat-stable extracellular hemolysin from Pseudomonas aeruginosa. Infection and Immunity 56:1385–1387.

49. Ochsner UA, Fiechter A, Reiser J. 1994. Isolation, characterization, and expression in Escherichia coli of the Pseudomonas aeruginosa rhlAB genes encoding a rhamnosyltransferase involved in rhamnolipid biosurfactant synthesis. Journal of Biological Chemistry 269:19787–19795.

50. Rahim R, Ochsner UA, Olvera C, Graninger M, Messner P, Lam JS, Soberón-Chávez G. 2001. Cloning and functional characterization of the Pseudomonas aeruginosa rhlC gene that encodes rhamnosyltransferase 2, an enzyme responsible for di-rhamnolipid biosynthesis. Molecular Microbiology 40:708–718.

51. Bazire A, Dufour A. 2014. The Pseudomonas aeruginosa rhlG and rhlAB genes are inversely regulated and RhlG is not required for rhamnolipid synthesis. BMC Microbiology 14:160.

52. Wittgens A, Tiso T, Arndt TT, Wenk P, Hemmerich J, Müller C, Wichmann R, Küpper B, Zwick M, Wilhelm S, Hausmann R, Syldatk C, Rosenau F, Blank LM. 2011. Growth independent rhamnolipid production from glucose using the non-pathogenic Pseudomonas putida KT2440. Microbial Cell Factories 10:80.

53. Chong H, Li Q. 2017. Microbial production of rhamnolipids: opportunities, challenges and strategies. Microbial Cell Factories 16:137.

54. Wilderman PJ, Vasil AI, Johnson Z, Wilson MJ, Cunliffe HE, Lamont IL, Vasil ML. 2001. Characterization of an Endoprotease (PrpL) Encoded by a PvdS-Regulated Gene in *Pseudomonas aeruginosa*. Infection and Immunity 69:5385–5394.

55. Zhang L, Tan FC, Strasfeld L, Hakki M, Kirienko NV. 2022. Long-Term Dominance of Carbapenem-Non-Susceptible Pseudomonas aeruginosa ST111 in Hematologic Malignancy Patients and Hematopoietic Cell Transplant Recipients. Frontiers in Cellular and Infection Microbiology 12.

56. Azghani AO, Miller EJ, Peterson BT. 2000. Virulence Factors from Pseudomonas aeruginosa Increase Lung Epithelial Permeability. Lung 178:261–269.

57. Zulianello L, Canard C, Köhler T, Caille D, Lacroix J-S, Meda P. 2006. Rhamnolipids Are Virulence Factors That Promote Early Infiltration of Primary Human Airway Epithelia by Pseudomonas aeruginosa. Infection and Immunity 74:3134–3147.

58. Vatsa P, Sanchez L, Clement C, Baillieul F, Dorey S. 2010. Rhamnolipid Biosurfactants as New Players in Animal and Plant Defense against Microbes. International Journal of Molecular Sciences 11:5095–5108.

59. Haba E, Pinazo A, Jauregui O, Espuny MJ, Infante MR, Manresa A. 2003. Physicochemical characterization and antimicrobial properties of rhamnolipids produced by Pseudomonas aeruginosa 47T2 NCBIM 40044. Biotechnology and Bioengineering 81:316–322.

60. Magalhães L, Nitschke M. 2013. Antimicrobial activity of rhamnolipids against Listeria monocytogenes and their synergistic interaction with nisin. Food Control 29:138–142.

61. Sha R, Meng Q. 2016. Antifungal activity of rhamnolipids against dimorphic fungi. The Journal of General and Applied Microbiology 62:233–239.

62. Marangon CA, Martins VCA, Ling MH, Melo CC, Plepis AMG, Meyer RL, Nitschke M. 2020. Combination of Rhamnolipid and Chitosan in Nanoparticles Boosts Their Antimicrobial Efficacy. ACS Applied Materials & Interfaces 12:5488–5499.

63. Gdaniec BG, Allard P-M, Queiroz EF, Wolfender J-L, van Delden C, Köhler T. 2020. Surface sensing triggers a broad-spectrum antimicrobial response in Pseudomonas aeruginosa. Environmental Microbiology 22:3572–3587.

64. Paulino BN, Pessôa MG, Mano MCR, Molina G, Neri-Numa IA, Pastore GM. 2016. Current status in biotechnological production and applications of glycolipid biosurfactants. Applied Microbiology and Biotechnology 100:10265–10293.

65. Davey ME, Caiazza NC, O’Toole GA. 2003. Rhamnolipid Surfactant Production Affects Biofilm Architecture in Pseudomonas aeruginosa PAO1. Journal of Bacteriology 185:1027–1036.

66. Jensen PØ, Bjarnsholt T, Phipps R, Rasmussen TB, Calum H, Christoffersen L, Moser C, Williams P, Pressler T, Givskov M, Høiby N. 2007. Rapid necrotic killing of polymorphonuclear leukocytes is caused by quorum-sensing-controlled production of rhamnolipid by Pseudomonas aeruginosa. Microbiology 153:1329–1338.

67. Köhler T, Guanella R, Carlet J, Delden Cv. 2010. Quorum sensing-dependent virulence during Pseudomonas aeruginosa colonisation and pneumonia in mechanically ventilated patients. Thorax 65:703–710.

68. Lee M-T. 2021. Micellization of Rhamnolipid Biosurfactants and Their Applications in Oil Recovery: Insights from Mesoscale Simulations. The Journal of Physical Chemistry B 125:9895–9909.

69. Esposito R, Speciale I, De Castro C, D’Errico G, Russo Krauss I. 2023. Rhamnolipid Self-Aggregation in Aqueous Media: A Long Journey toward the Definition of Structure– Property Relationships. International Journal of Molecular Sciences 24:5395.

70. Gdaniec BG, Bonini F, Prodon F, Braschler T, Köhler T, van Delden C. 2022. Pseudomonas aeruginosa rhamnolipid micelles deliver toxic metabolites and antibiotics into Staphylococcus aureus. iScience 25:103669.

71. Kownatzki R, Tümmler B, Döring G. 1987. RHAMNOLIPID OF PSEUDOMONAS AERUGINOSA IN SPUTUM OF CYSTIC FIBROSIS PATIENTS. The Lancet 329:1026–1027.

72. Bjarnsholt T, Jensen PØ, Jakobsen TH, Phipps R, Nielsen AK, Rybtke MT, Tolker-Nielsen T, Givskov M, Høiby N, Ciofu O, Consortium tSCFS. 2010. Quorum Sensing and Virulence of Pseudomonas aeruginosa during Lung Infection of Cystic Fibrosis Patients. PLOS ONE 5:e10115.

73. Bhagirath AY, Li Y, Somayajula D, Dadashi M, Badr S, Duan K. 2016. Cystic fibrosis lung environment and Pseudomonas aeruginosa infection. BMC Pulmonary Medicine 16.

74. Faure E, Kwong K, Nguyen D. 2018. Pseudomonas aeruginosa in Chronic Lung Infections: How to Adapt Within the Host? Frontiers in immunology 9.

75. Liberati NT, Urbach JM, Miyata S, Lee DG, Drenkard E, Wu G, Villanueva J, Wei T, Ausubel FM. 2006. An ordered, nonredundant library of Pseudomonas aeruginosa strain PA14 transposon insertion mutants. Proceedings of the National Academy of Sciences of the United States of America 103:2833–2838.

76. Jacobs MA, Alwood A, Thaipisuttikul I, Spencer D, Haugen E, Ernst S, Will O, Kaul R, Raymond C, Levy R, Chun-Rong L, Guenthner D, Bovee D, Olson MV, Manoil C. 2003. Comprehensive transposon mutant library of Pseudomonas aeruginosa. Proceedings of the National Academy of Sciences 100:14339–14344.

77. Held K, Ramage E, Jacobs M, Gallagher L, Manoil C. 2012. Sequence-Verified Two-Allele Transposon Mutant Library for Pseudomonas aeruginosa PAO1. Journal of Bacteriology 194:6387–6389.

78. Zhang L, Gade V, Kirienko NV. 2023. Pathogen-induced dormancy in liquid limits gastrointestinal colonization of *Caenorhabditis elegans*. Virulence 14.

