## Supplementary material for "Cytotoxic rhamnolipid micelles drive acute virulence in *Pseudomonas aeruginosa*": Sup Figures and Tables

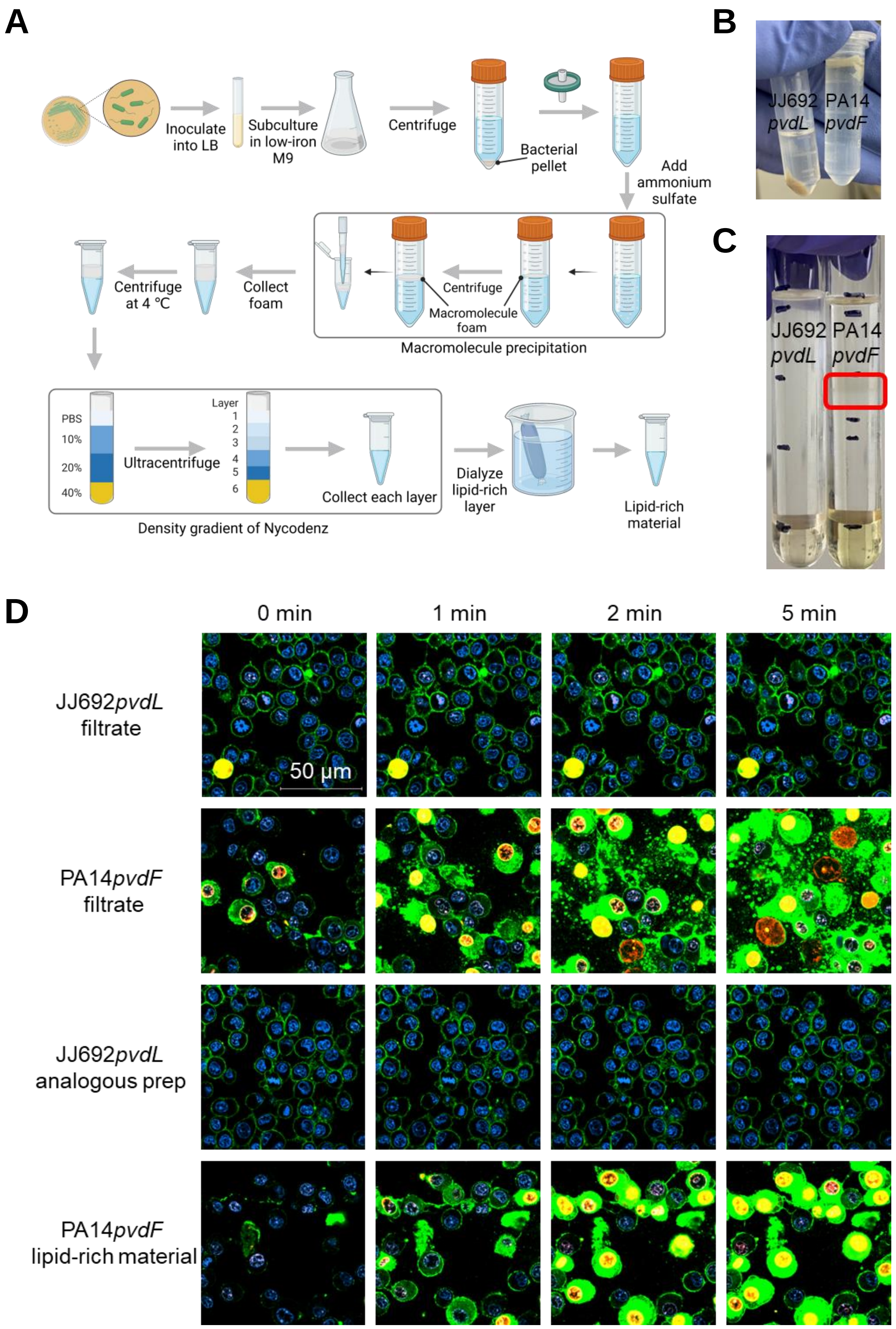
A
Inoculate into LB
Subculture in low-iron M9
Centrifuge
Bacterial pellet
Add ammonium sulfate
Centrifuge
Macromolecule foam
Macromolecule precipitation
Collect foam
Centrifuge at 4 °C
PBS
10%
20%
40%
Ultracentrifuge
Layer
1
2
3
4
5
6
Collect each layer
Density gradient of Nycodenz
Dialyze lipid-rich layer
Lipid-rich material
B
JJ692 pvdL
PA14 pvdF
C
JJ692 pvdL
PA14 pvdF
D
0 min
1 min
2 min
5 min
JJ692pvdL filtrate
50 µm
PA14pvdF filtrate
JJ692pvdL analogous prep
PA14pvdF lipid-rich material

### Fig. S1

**Fig. S1 Purified lipid-rich material from *P. aeruginosa* is cytotoxic to murine macrophages. (A)** Detailed schematic of purification pipeline for *P. aeruginosa* lipid-rich material. Created with BioRender. **(B)** Representative image of macromolecule floc (right, PA14*pvdF*) and pellet (left, JJ692*pvdL*) after ammonium sulfate precipitation. **(C)** Representative image of Nycodenz density gradient after ultracentrifugation for PA14*pvdF* (right – layer with high FM 1-43 fluorescence highlighted in red) and JJ692*pvdL* (left). **(D)** Interactions between RAW264.7 cells and bacterial filtrate or purified lipid-rich material from PA14*pvdF* or material from JJ692*pvdL* in the presence of SYTOX Orange cell-impermeant nucleic acid stain [red]. Secreted bacterial lipids were prelabeled with FM 1-43 [green]. Cells were prelabeled with Hoechst 33342 cell-permeant nucleic acid stain [blue].

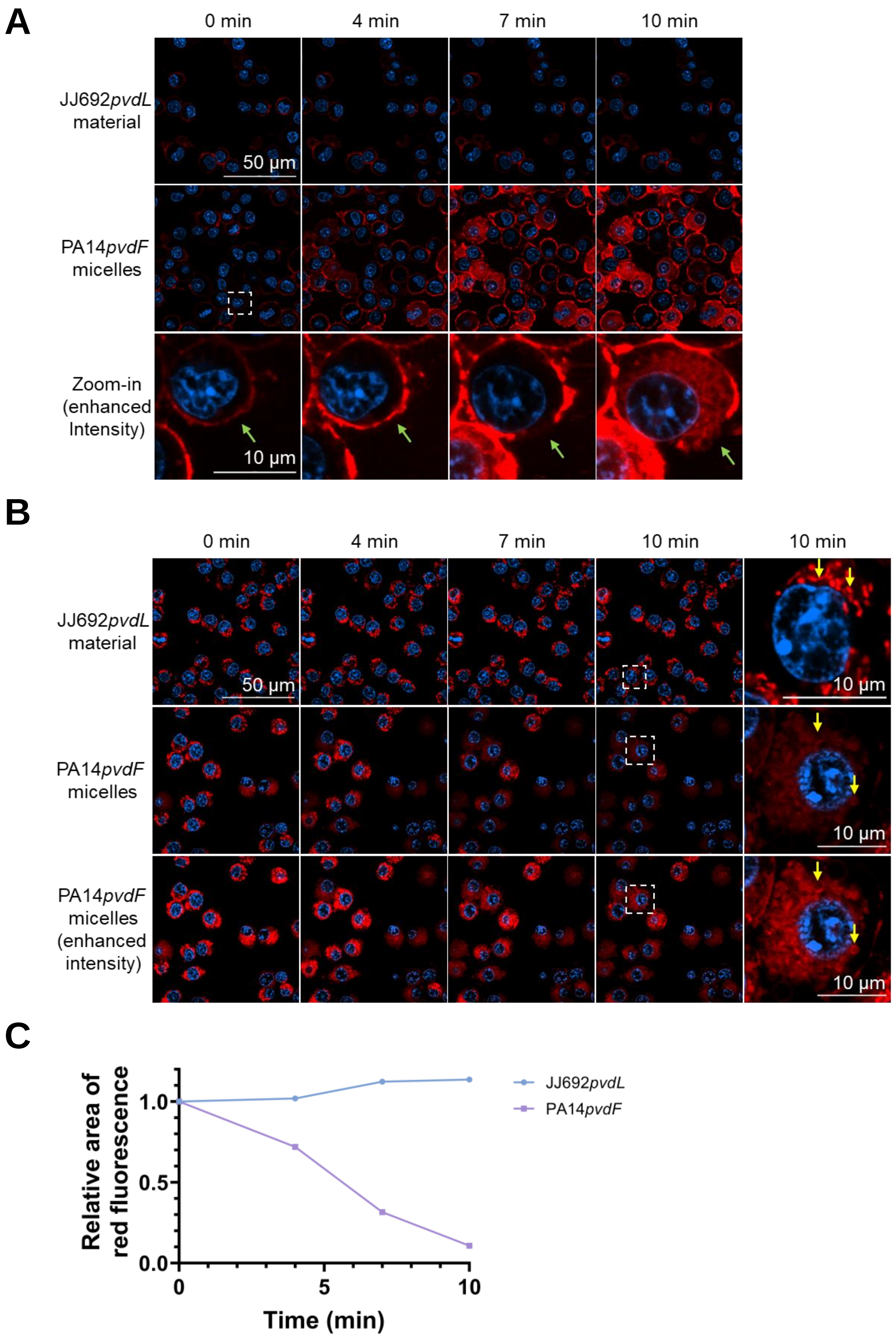
A
0 min
4 min
7 min
10 min
JJ692pvdL material
50 µm
PA14pvdF micelles
Zoom-in (enhanced Intensity)
10 µm
B
0 min
4 min
7 min
10 min
10 min
JJ692pvdL material
50 µm
10 µm
PA14pvdF micelles
10 µm
PA14pvdF micelles (enhanced intensity)
10 µm
C
Relative area of red fluorescence
1.0
0.5
0.0
0
5
10
Time (min)
JJ692pvdL
PA14pvdF

### Fig. S2

**Fig. S2 Purified *P. aeruginosa* micelles rupture plasma membrane and damage mitochondrial membranes in murine macrophages. (A)** Visualization of macrophage plasma membrane after 10 min exposure to purified micelles from PA14*pvdF* or material from JJ692*pvdL*. A representative cell (white square) was selected and enhanced for detailed view of the plasma membrane (green arrow). Cells were prelabeled with Hoechst 33342 [blue] and CellMask Deep Red plasma membrane stain [red]. **(B)** Visualization of macrophage mitochondria after 10 min exposure to purified micelles from PA14*pvdF* or material from JJ692*pvdL*. A representative cell (white square) was selected and enhanced for detailed view of individual mitochondria (yellow arrow). Cells were pre-labeled with Hoechst 33342 [blue] and MitoTracker Red CMXRos [red]. **(C)** Change in area of red fluorescence from stained mitochondria after micelle exposure. Images were analyzed via Fiji. The area with MitoTracker Red CMXRos fluorescence at each time point was normalized to the one at 0 min.

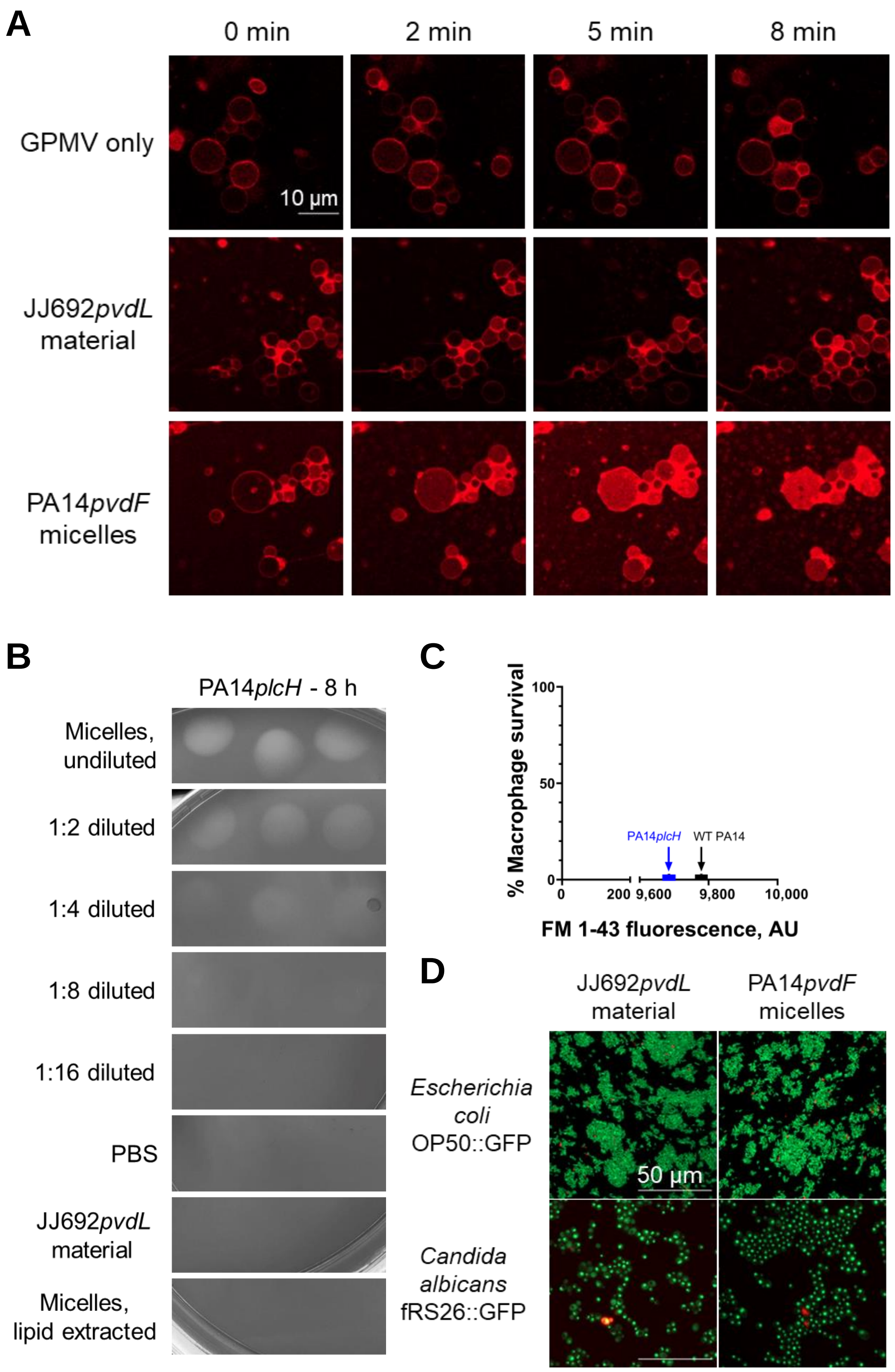
A
0 min
2 min
5 min
8 min
GPMV only
10 µm
JJ692*pvdL* material
PA14*pvdF* micelles
B
PA14*plcH* - 8 h
Micelles, undiluted
1:2 diluted
1:4 diluted
1:8 diluted
1:16 diluted
PBS
JJ692*pvdL* material
Micelles, lipid extracted
C
% Macrophage survival
100
50
0
PA14*plcH*
WT PA14
0
200
9,600
9,800
10,000
FM 1-43 fluorescence, AU
D
JJ692*pvdL* material
PA14*pvdF* micelles
*Escherichia coli* OP50::GFP
50 µm
*Candida albicans* fRS26::GFP

### Fig. S3

**Fig. S3 *P. aeruginosa* micelles kill eukaryotic and prokaryotic cells. (A)** Visualization of giant plasma membrane vesicles (GPMVs) derived from human bronchial epithelial cells (16HBE) after 8 min exposure to purified micelles from PA14*pvdF* or material from JJ692*pvdL*. GPMVs were prelabeled with CellMask Deep Red plasma membrane stain. **(B)** Hemolysis of erythrocytes on sheep blood's agar after 8 h exposure to purified micelles from PA14*plcH*, material from JJ692*pvdL*, or PA14*plcH* sample after lipid extraction via chloroform. **(C)** Murine macrophage (RAW264.7) survival after exposure to supernatants from wild-type PA14 and PA14*plcH* (grown in low-iron M9 medium). Survival was normalized to saline control. **(D)** Visualization of *Escherichia coli* OP50::GFP and *Candida albicans* fRS26::GFP after 4 h exposure to purified micelles from PA14*pvdF* or material from JJ692*pvdL*.

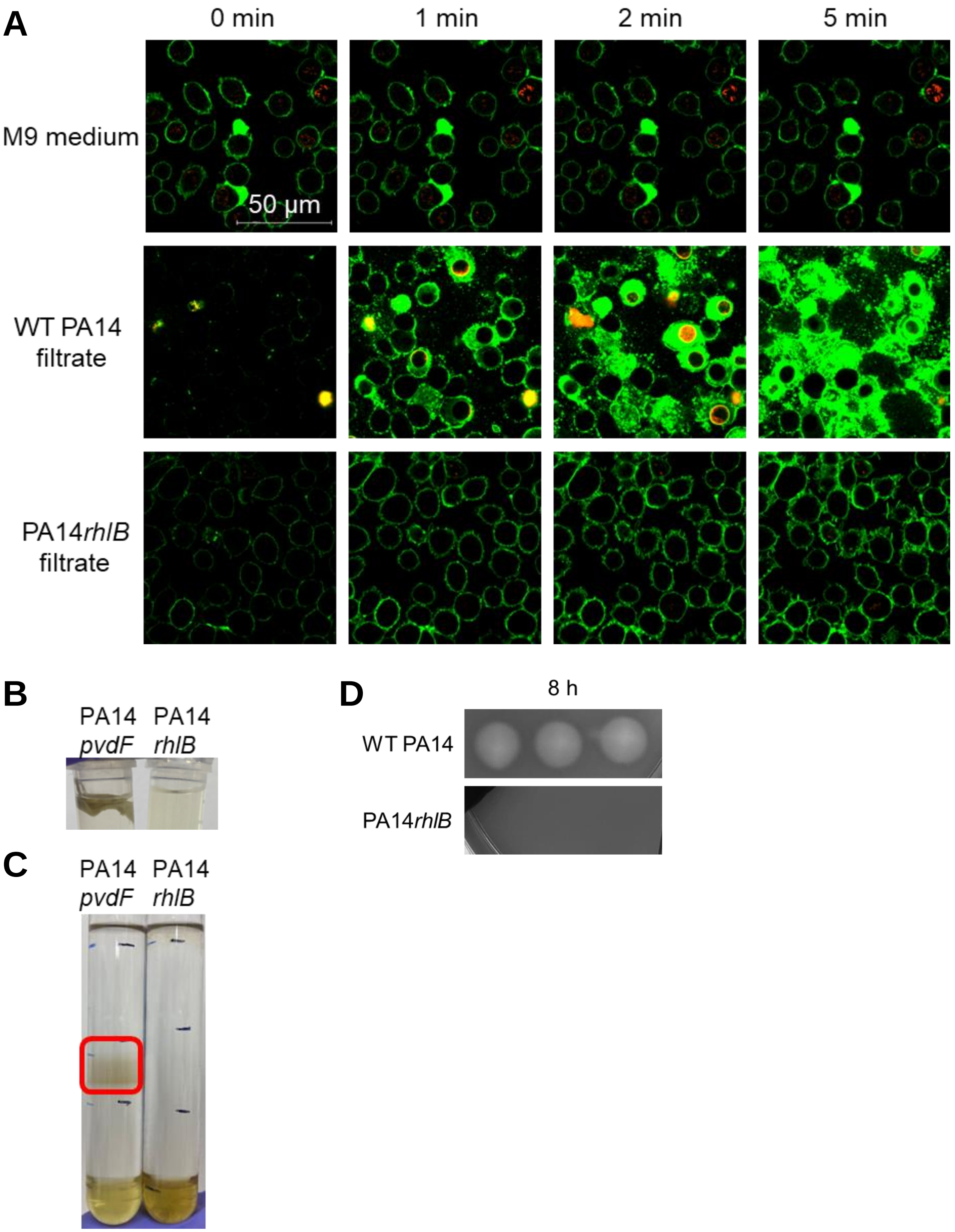
A
0 min
1 min
2 min
5 min
M9 medium
50 µm
WT PA14 filtrate
PA14rhlB filtrate
B
PA14 pvdF
PA14 rhlB
C
PA14 pvdF
PA14 rhlB
D
8 h
WT PA14
PA14rhlB

### Fig. S4

**Fig. S4 Rhamnolipid biosynthetic mutants do not produce cytotoxic micelles. (A)** Interactions between RAW264.7 cells and bacterial filtrate from wild-type PA14 or PA14*rhlB* in the presence of SYTOX Orange cell-impermeant nucleic acid stain [red]. Secreted bacterial lipids were prelabeled with FM 1-43 [green]. **(B)** Representative image of macromolecule floc (right, PA14*pvdF*) after ammonium sulfate precipitation. No floc was formed in PA14*rhlB* filtrate with ammonium sulfate. **(C)** Representative image of Nycodenz density gradient after ultracentrifugation for PA14*pvdF* (left – layer with high FM 1-43 fluorescence highlighted in red) and PA14*rhlB* (right). **(D)** Hemolysis of erythrocytes on sheep blood's agar after 8 h exposure to purified micelles from wild-type PA14 and material from PA14*rhlB*.

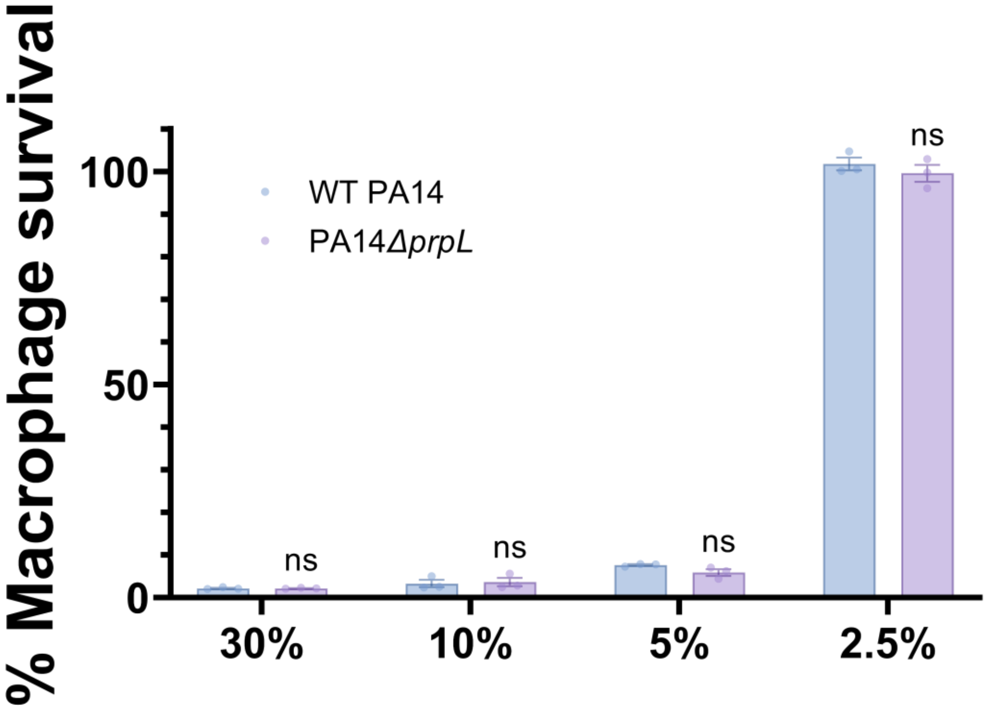


**Fig. S5 Micelle cytotoxicity was independent of proteins.** Cytotoxicity of bacterial filtrate from wild-type PA14 and PA14Δ*prpL* against RAW264.7 cells. Error bars represent SEM of 3 biological replicates. ns, not statistically significant based on two-way ANOVA.

Fig. S5

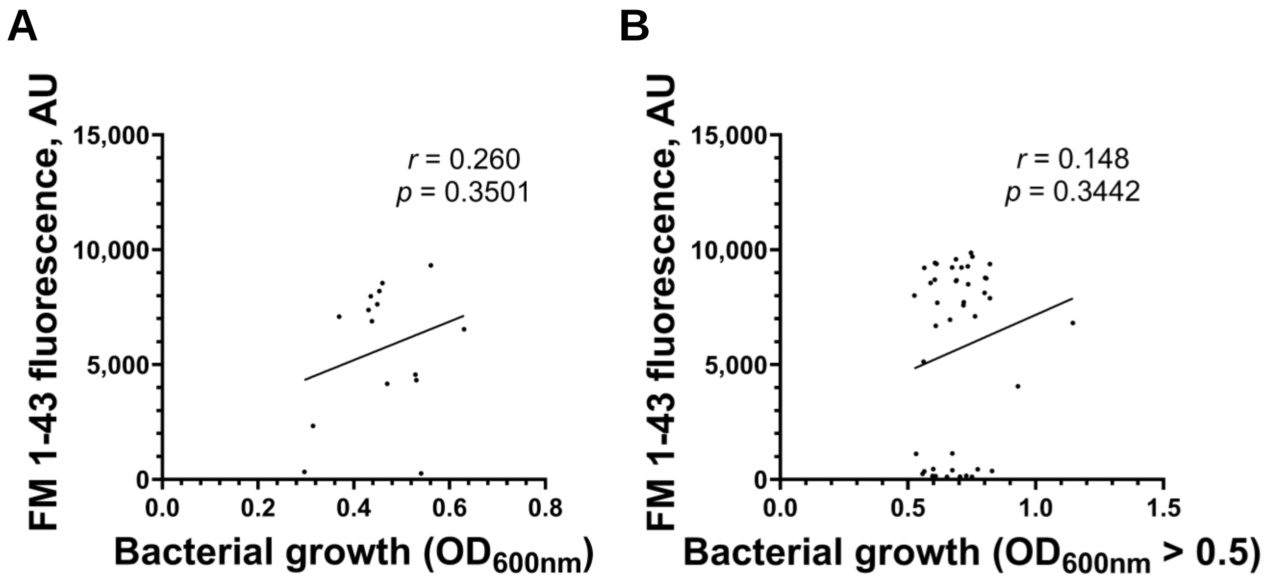


**Fig. S6 There is no correlation between bacterial growth and supernatant rhamnolipid content in *P. aeruginosa* clinical isolates. (A)** Correlation between bacterial growth ($OD_{600nm}$) in the overnight cultures of 12 hematological isolates and FM 1-43 fluorescence in the supernatants. **(B).** Correlation between bacterial growth ($OD_{600nm}$) in the overnight cultures of pediatric cystic fibrosis clinical isolates and FM 1-43 fluorescence in the supernatants. From the full panel of 68 isolates, we removed those with severely poor bacterial growth ($OD_{600nm} < 0.5$).

### Fig. S6

**Movie S1 Interactions between RAW264.7 cells and material from JJ692*pvdL*.** The material was prelabeled with FM 1-43 [green]. Cells prelabeled with Hoechst 33342 cell-permeant nucleic acid stain [blue], were in the serum-free medium containing SYTOX Orange cell-impermeant nucleic acid stain [red]. Images were taken every 30 seconds. 2-second video corresponds to 5-minute imaging, showing 5 frames per second. Scale bar = 50 µm.

**Movie S2 Interactions between RAW264.7 cells and purified lipid-rich material from PA14*pvdF*.** Secreted bacterial lipids were prelabeled with FM 1-43 [green]. Cells prelabeled with Hoechst 33342 cell-permeant nucleic acid stain [blue], were in the serum-free medium containing SYTOX Orange cell-impermeant nucleic acid stain [red]. Images were taken every 30 seconds. 2-second video corresponds to 5-minute imaging, showing 5 frames per second. Scale bar = 50 µm.

**Table S1** Phospholipids detected in liquid chromatography-mass spectrometry (LC-MS).

| Label | Mass | m/z | Vol % |
|---|---|---|---|
| **Glycerophosphocholines (PC) (major membrane component)** | | | |
| PC not found | | | |
| **Phosphatidylethanolamines (PE) (major membrane component)** | | | |
| PE(6:0/6:0)[U]; C17 H34 N O8 P | 411.2 | 410.19 | 0.01 |
| PE(8:0/8:0)[U]; C21 H42 N O8 P | 467.26 | 466.26 | 0 |
| **Glycerophosphoglycerols (PG) (major membrane component)** | | | |
| PG(10:0/10:0); C26 H51 O10 P | 554.32 | 553.31 | 0.02 |
| **Glycerophosphoinositols (PI)** | | | |
| PI(18:4(6Z,9Z,12Z,15Z)/12:0); C39 H67 O13 P | 774.44 | 773.43 | 0.02 |
| PI(22:2(13Z,16Z)/18:1(9Z)); C49 H89 O13 P | 916.61 | 915.6 | 0 |
| **Glycerophosphoserines (PS)** | | | |
| PS(13:0/0:0); C19 H38 N O9 P | 455.23 | 454.22 | 0.01 |
| PS(17:0/0:0); C23 H46 N O9 P | 511.29 | 510.28 | 0 |
| **Overall** | | | 0.06 |

### Table S1

**Table S2** Proteins detected in liquid chromatography-mass spectrometry (LC-MS).

| | Score | Mass (Da) | Matches | Match (sig) | Sequences | Seq (sig) | emPAI | Description |
|---|---|---|---|---|---|---|---|---|
| 1 | 353 | 48223 | 35 | 19 | 5 | 4 | 0.47 | Protease IV Tax= *Pseudomonas aeruginosa* |
| 2 | 166 | 24394 | 46 | 7 | 5 | 3 | 0.77 | sp\|TRYP_PIG\| |
| 3 | 68 | 57526 | 10 | 3 | 3 | 1 | 0.08 | Cypermethrin hydrolyzing aminopeptidase Tax= *Pseudomonas aeruginosa* |
| 4 | 48 | 50533 | 13 | 1 | 3 | 1 | 0.1 | Neutral metalloproteinase (Fragment) Tax= *Pseudomonas aeruginosa* |
| 5 | 29 | 111839 | 6 | 1 | 1 | 1 | 0.04 | amino acid adenylation domain-containing protein Tax= *Micromonospora sp.* ANENR4 |
| 6 | 29 | 37088 | 1 | 1 | 1 | 1 | 0.13 | SIS domain-containing protein Tax= *Actinobacteria bacterium* |

### Table S2

**Table S3** Sequences of rhamnolipid biosynthetic enzymes and known quorum-sensing regulators in *P. aeruginosa* isolates compared to the reference strain PAO1.

| Strain | FM1-43 | Vfr | LasR | RsaL | LasI | RhlI | RhlR | RhlA | RhlB | RmlA | RmlB | RmlC | RmlD | Strain Source |
|---|---|---|---|---|---|---|---|---|---|---|---|---|---|---|
| JJ692 | 311.5 | - | D65G | - | - | S62G | - | Q168P | - | E165K | - | - | V111A, C201R, V204A, A253T, R272H | 39 |
| U2504 | 509 | - | G235S | - | - | S62G | Y234* | Q168P | - | E165K | - | - | V111A, C201R, V204A, A253T, R272H | 39 |
| E2 | 616.5 | - | - | - | - | S62G, D83E | - | Q168P | D70N | E165K | Q73P | - | V111A, C201R, V204A | 39 |
| X13273 | 401 | - | - | - | - | S62G, D83E | Δ104bp | Q168P | D382N | E165K, Q211H, E290G | - | - | V111A, C201R, V204A, A253T | 39 |
| X24509 | 251 | - | Δ14bp (114-127/720nt) | - | - | D83E | +4bp (703/726nt) | Q168P | - | - | - | V146L | A91G, C201R | 39 |
| CF27 | 363 | - | - | - | - | S62G, D83E | - | Q168P | - | D188N | E71D | - | C201R, V204A | 39 |
| CF5 | 139.5 | - | A27V, R180Q | - | - | S62G, D83E | - | Q168P | S251N | - | E71D | Q96H | A91G, C201R | 39 |
| PA3-17 | 693 | - | - | - | Q57* | D83E | Q25* | Q168P | - | - | - | A173T | C201R, V204A | 34 |
| PA3-22 | 782 | H164R | L128P | - | - | S62G, D83E | - | Q168P | - | - | - | A173T | C201R, V204A | 34 |
| M0134 | 328.33 | - | - | - | - | S62G, D83E | A232D | Q168P | - | D188N | - | - | A91G, V111A, C201R | 55 |
| PA2-59 | 1893.5 | - | - | - | - | S62G, D83E | D98G | Q168P | - | - | E71D | - | A91G, S110N, V111A, C201R, V204A | 34 |
| PA2-94 | 1377.5 | - | - | - | - | S62G, D83E | - | Q168P | - | D188N | - | - | C201R, V204A | 34 |
| PA3-25 | 3086.5 | - | - | - | - | S62G, D83E | - | Q168P | - | D188N | - | - | C201R, V204A | 34 |
| PAHP3 | 2634.5 | - | Δ7bp (676-682/720nt) | - | - | S62G, D83E | - | Q168P | - | R209H, E290G | - | - | A196V, C201R, R272H | 34 |
| M0101 | 2329 | - | W152* | - | - | S62G, D83E | - | Q168P | T23S | D188N | - | - | A91G, V111A, C201R | 55 |
| PA14 | 9622 | - | - | - | - | S62G, A127G | - | Q168P | - | E165K | - | - | V111A, C201R, V204A, R272H | 39 |
| 6077 | 8639 | - | V226I | - | - | S62G | - | Q168P | - | E165K | - | - | V111A, C201R, V204A, A253T, R272H | 39 |
| S35004 | 7066.5 | - | K16* | - | - | S62G, D83E | - | Q168P | I36V, A81T | - | E71D | Q96H | A196V, C201R, V204A | 39 |
| S54485 | 6109.5 | - | - | - | - | S62G, D83E | - | Q168P | D382N | E165K, Q211H, E290G | - | - | V111A, C201R, V204A, A253T | 39 |
| 62 | 8745 | - | +2bp (417/720nt) | - | - | D83E | - | Q168P | - | - | E71D | - | V47I, A91G, V111A, C201R, V204A | 39 |
| MSH3 | 8611.5 | - | - | - | - | S62G, D83E | - | Q168P. H240R | - | N101D, E165K, E290G | - | - | D72N, A91G, C201R, V204A, R272H | 39 |
| MSH10 | 8839 | - | - | - | - | S62G, D83E | - | Q168P, H240R | - | N101D, E165K, E290G | - | - | D72N, A91G, C201R, V204A, R272H | 39 |
| CF18 | 9358 | - | - | - | - | S62G, D83E | - | Q168P | - | E290G | E71D | - | A91G, C201R | 39 |
| PA2-61 | 10209 | - | - | - | - | S62G, D83E | - | Q168P | - | E290G | E71K | Q96H, A173S | V111A, A135T, C201R, V204A | 34 |
| PA2-87 | 9452 | +1bp (444/645nt) | - | - | - | S62G, D83E | - | Q168P | - | - | - | V146L | C201R, V204A | 34 |
| PA2-72 | 9148 | - | - | - | - | S62G, D83E | - | Q168P | T62A | - | - | Q96H | A91G, C201R | 34 |
| PA2-45 | 11413 | - | R224H | - | - | S62G | - | Q168P | - | E165K | - | - | V111A, C201R, V204A, A253T, R272H | 34 |
| PA5-40 | 9096 | T5A | - | - | - | D83E | - | Q168P | S251N | - | E71D | - | C201R, L282M | 34 |
| M0087 | 7376.67 | - | W152* | - | - | S62G, D83E | - | Q168P | T23S | D188N | - | - | A91G, V111A, C201R | 55 |
| M0177 | 7086.33 | - | W152* | - | - | S62G, D83E | - | Q168P | T23S | D188N | - | - | A91G, V111A, C201R | 55 |
| M0128 | 8546.33 | - | - | - | - | D83E | - | Q168P | S251N | - | E71D | - | C201R, L282M | 55 |
| M0162 | 9318 | - | - | - | - | D83E | - | Q168P | - | D188N | - | - | C201R, V204A | 55 |
| M0068 | 6539.67 | - | T115I | - | - | S62G, D83E | - | Q168P | S251N | D188N | - | - | A91G, C201R | 55 |

### Table S3
